## Supplementary Information for "Rare and common vertebrates span a wide spectrum of population trends"

**This PDF file includes:**

- Tables S1 to S8
- Figures S1 to S19

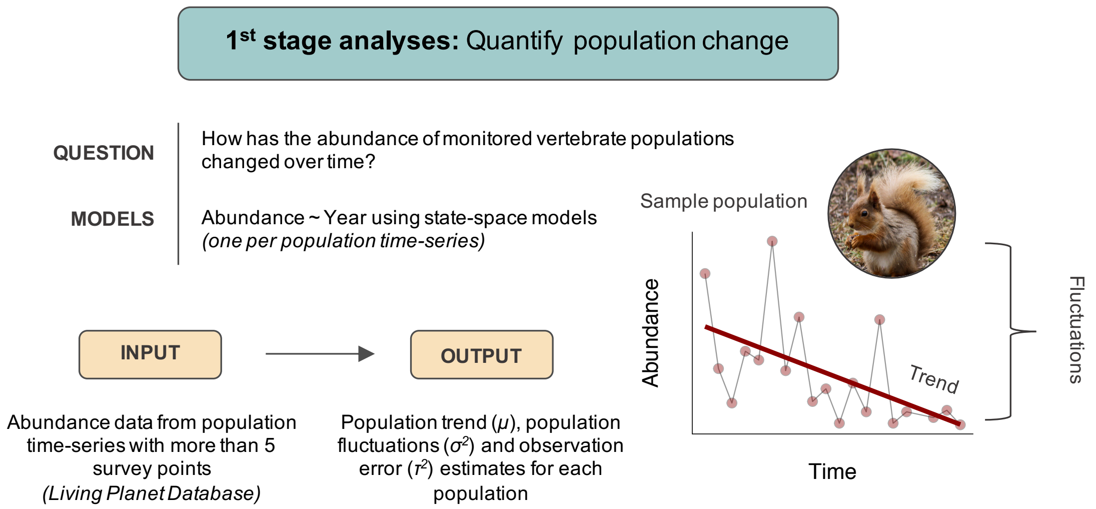


**Figure S1. Conceptual diagram of the first stage of our analyses where we calculated population trends and fluctuations.** We analysed vertebrate population time-series from the Living Planet Database (133,092 records) covering the period between 1970 and 2014. These time-series represent repeated monitoring surveys of the number of individuals in a given area (species' abundance over time), to which we refer as "populations". Diagram shows one sample population of Red squirrel (*Sciurus vulgaris*). We quantified two aspects of population change – overall change in abundance over time (population trends) and abundance variability over time (population fluctuations). We used state-space models that account for observation error and random fluctuations[1]. The input abundance data for the state-space models were scaled to a common magnitude between zero and one to analyse within-population relationships to prevent conflating within-population relationships and between-population relationships[2]. See methods for additional details.

### **1st stage analyses:** Quantify population change

**QUESTION** How has the abundance of monitored vertebrate populations changed over time?

**MODELS** Abundance ~ Year using state-space models *(one per population time-series)*

**INPUT** → **OUTPUT**

Abundance data from population time-series with more than 5 survey points *(Living Planet Database)*

Population trend ($\mu$), population fluctuations ($\sigma^2$) and observation error ($\tau^2$) estimates for each population

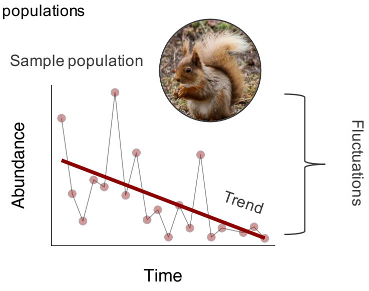


### **2nd stage analyses:** Test heterogeneity in population trends and fluctuations

**Prior structure 1:**
Hierarchical models in a Bayesian framework with weakly informative (flat) priors

*Pr(μ) ~ N(0, 10⁸)*
*Pr(σ²) ~ Inverse Wishart (V = 0, nu = 0)*

**Prior structure 2:**
Hierarchical models in a Bayesian framework with weakly informative (parameter expanded) priors and a variance-covariance structure that allows the slopes of population trends and fluctuations to covary for each random effect.

*Pr(μ) ~ N(0, 10⁸)*
*Pr(σ²) ~ Inverse Wishart (V = 1, nu = 1)*

**INPUT**

Population trend ($\mu$), population fluctuations ($\sigma^2$) and observation error ($\tau^2$) estimates for each population

**GEOGRAPHIC PATTERNS**

**QUESTIONS** How do vertebrate population trends and fluctuations vary across latitudes, realms and biomes?

**MODELS**
*μ ~ latitude, random = species*
*μ ~ 1 + realm, random = species*
*μ ~ 1 + biome, random = species*
*σ² ~ latitude, random = species*
*σ² ~ 1 + realm, random = species*
*σ² ~ 1 + biome, random = species*

**+**

Bird, amphibian and reptile phylogenies

**TAXONOMIC PATTERNS**

**QUESTIONS** How do vertebrate population trends and fluctuations vary across taxa and phylogenies?

**MODELS**
*μ ~ 1 + taxa, random = species*
*σ² ~ 1 + taxa, random = species*
*μ ~ 1, random = species + phylogeny*
*σ² ~ 1, random = species + phylogeny*

**Prior structure 2** (phylogeny models); **Prior structure 1** (geographic and taxonomic models)

**+**

Bird and mammal geographic range, species' mean population size, species' habitat specificity

**RARITY PATTERNS**

**QUESTIONS** How do vertebrate population trends and fluctuations vary across rarity metrics?

**MODELS**
*μ ~ log(geographic range), random = species*
*μ ~ log(mean population size), random = species*
*μ ~ habitat specificity, random = species*
*σ² ~ log(geographic range), random = species*
*σ² ~ log(mean population size), random = species*
*σ² ~ habitat specificity, random = species*

**+**

Species' IUCN conservation status and IUCN threats categories

**IUCN CONSERVATION STATUS AND TREATS**

**QUESTIONS** How do vertebrate population trends and fluctuations vary across species' IUCN Red List Categories and type and number of threats?

*μ ~ 1 + IUCN Red List Category, random = species*
*μ ~ 1 + threat type, random = species*
*μ ~ number of threats, random = species*
*σ² ~ 1 + IUCN Red List Category, random = species*
*σ² ~ 1 + threat type, random = species*
*σ² ~ number of threats, random = species*

**Prior structure 1** (rarity and IUCN models)

**OUTPUT**

Effect sizes for each tested metric

34

**Figure S2. Conceptual diagram of the second stage of our analyses where we quantified the geographic, taxonomic, rarity and threat patterns within vertebrate population trends and fluctuations.** We modelled the trend and fluctuation estimates from the first stage (Figure S1) across latitude, realm, biome, taxa, rarity metrics, phylogenetic relatedness, species' conservation status and threat type using a Bayesian modelling framework[3]. Each model included a species random intercept effect to account for the possible correlation between the trends of populations from the same species. The prior structure (weakly informative priors) was identical across all models except the phylogeny models from the taxonomic patterns section, where the prior structure allowed for an additional phylogeny random effect. See methods for additional details.

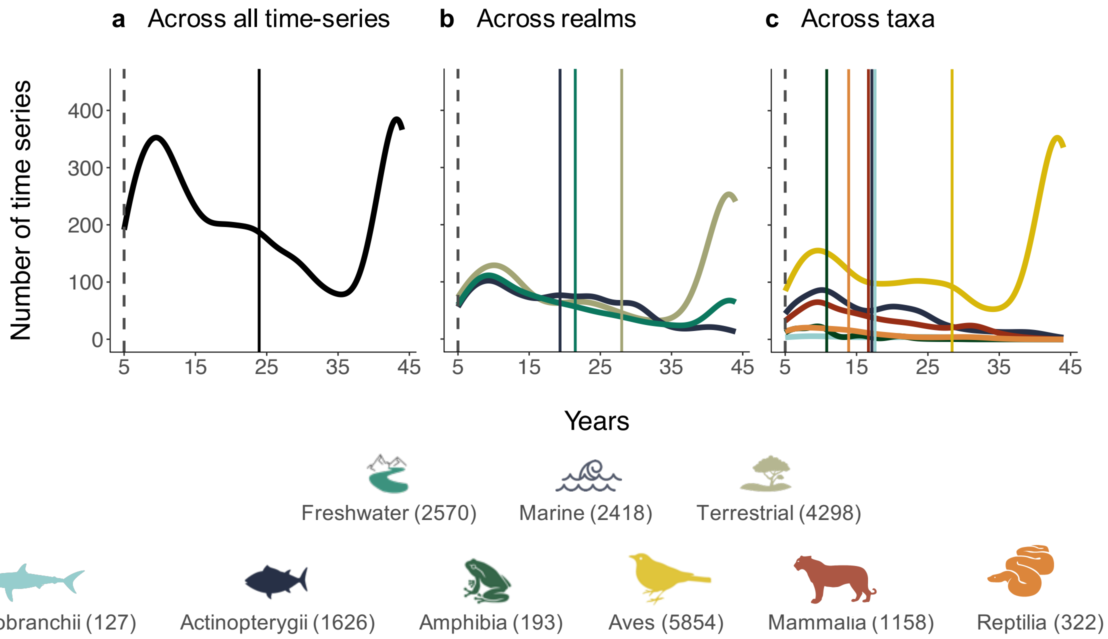


**Figure S3. The duration of monitoring varies by realm and taxa.** Distribution of monitoring duration across (a) all time-series, (b) realms and (c) taxa. In our study, we included time-series with more than five survey points in time, with the dashed line representing five years and solid lines showing the mean duration for each category. Numbers in legend correspond to sample size in each category.

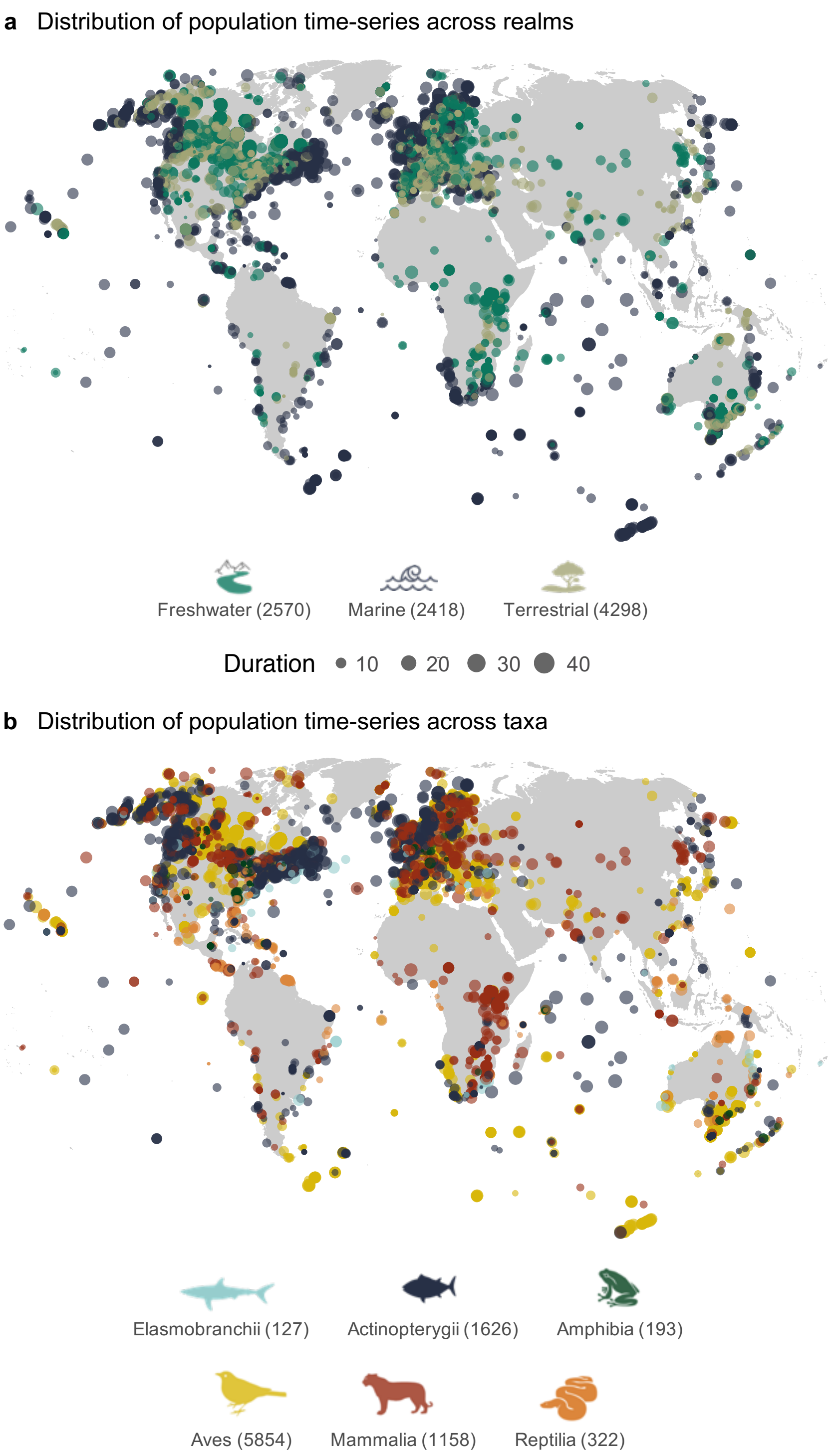


**Figure S5. The Living Planet Data represents a broad range of geographic locations, ecological settings and taxonomic groups.** Our analysis of the patterns in vertebrate population trends and fluctuations includes time-series across realms (a) and different taxa (b), with a global geographic distribution of records. Numbers in legend correspond to sample size in each category.

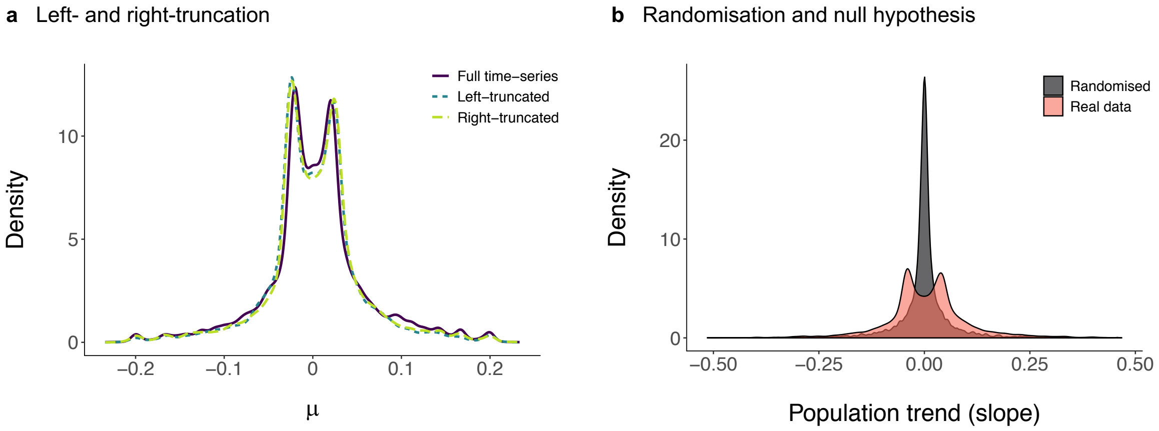


**Figure S6. The distribution of population trend values across time-series was not sensitive to the omission of the first five (left-truncation) or the last five years (right-truncation) of population records and it differed from a null distribution derived from randomised data.** Following Fournier *et al.* 2019[4], we tested the time-series that we analysed for site-selection bias. Removing the first five survey points reduces the bias stemming from starting population surveys at points when individual density is high, whereas removing the last five years reduces the bias of starting surveys when species are very rare. There were slightly fewer trends centred on zero (no net change in abundance over time) when we left- and right-truncated the data, suggesting that longer time-series are more likely to show no net changes in abundance (see Figure S7 for a visualization of population trends versus monitoring duration. We also compared the distribution of estimated population trends against a null hypothesis (b). To derive a null distribution, we used a randomisation approach. Within each time series, we randomised the abundance data, keeping the overall range of the original data. The two peaks of $\mu$ are apparent in the overall distribution of time series data. These peaks are created by many weakly positive and negative population trends from longer time series that often are bird species from terrestrial systems. We hypothesise that there might a publication bias against no net change studies, or a bias against including such studies in global databases.

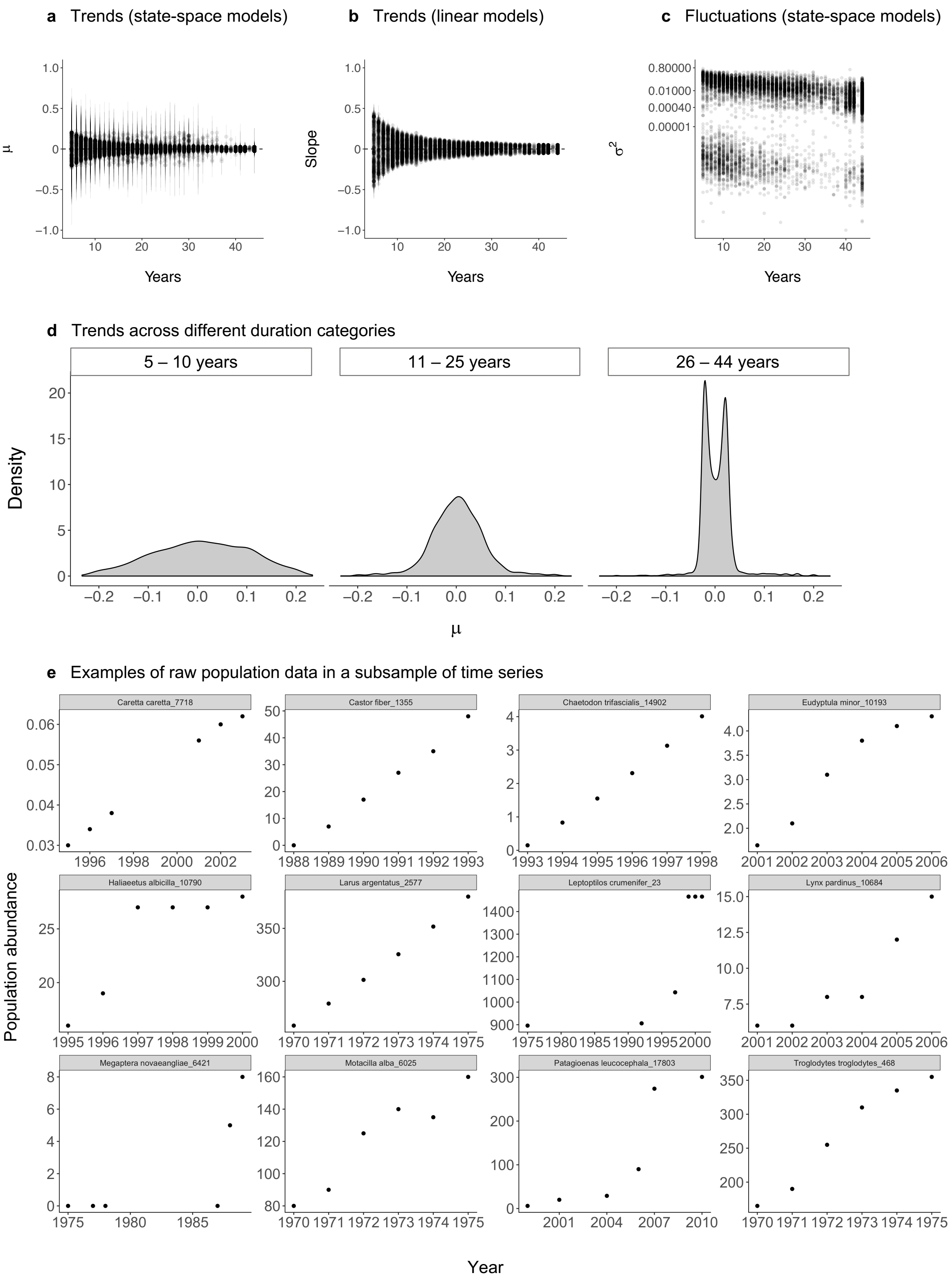


**Figure S7. Both positive and negative vertebrate population trends were smaller in magnitude for longer time-series of data.** Monitoring duration results are for 9286 populations from 2084 species. Population trends ($\mu$) were estimated for all populations monitored for more than five time points using state-space models (a, d) and linear models (b). Population fluctuations (c) are plotted on a log10 y axis and represent the estimates for process noise ($\sigma^2$, the process noise is the total variance around the population trend minus the variance attributed to observation error) derived from state-space models. Error bars on (a) and (b) show 95% confidence intervals. The sample sizes for the duration categories were as follows 5 - 10 years: 2084 time series; 10 - 25 years: 3358 time series; 25 - 44 years; 3844 time series. Plot (e) shows the raw population trend data behind 12 time series which had the same population trend values ($\mu = 0.20$). These time series are part of a “band” of time series which had very similar population trend estimates. Eighty, or approximately 1% of the time series we analysed form linear relationships over time with errors around the slopes of $<0.001$, such that we suspect these data might be modelled rather than measured population data. The presence of modelled data within the dataset may help partially explain the low variance bands of $\sigma^2$ values (c) and the pattern of two peaks in weak population increases and decreases for longer time series (d). Please see Methods sections “Time series with low variation” and “Clustering in the values of population trends and fluctuations” for further details.

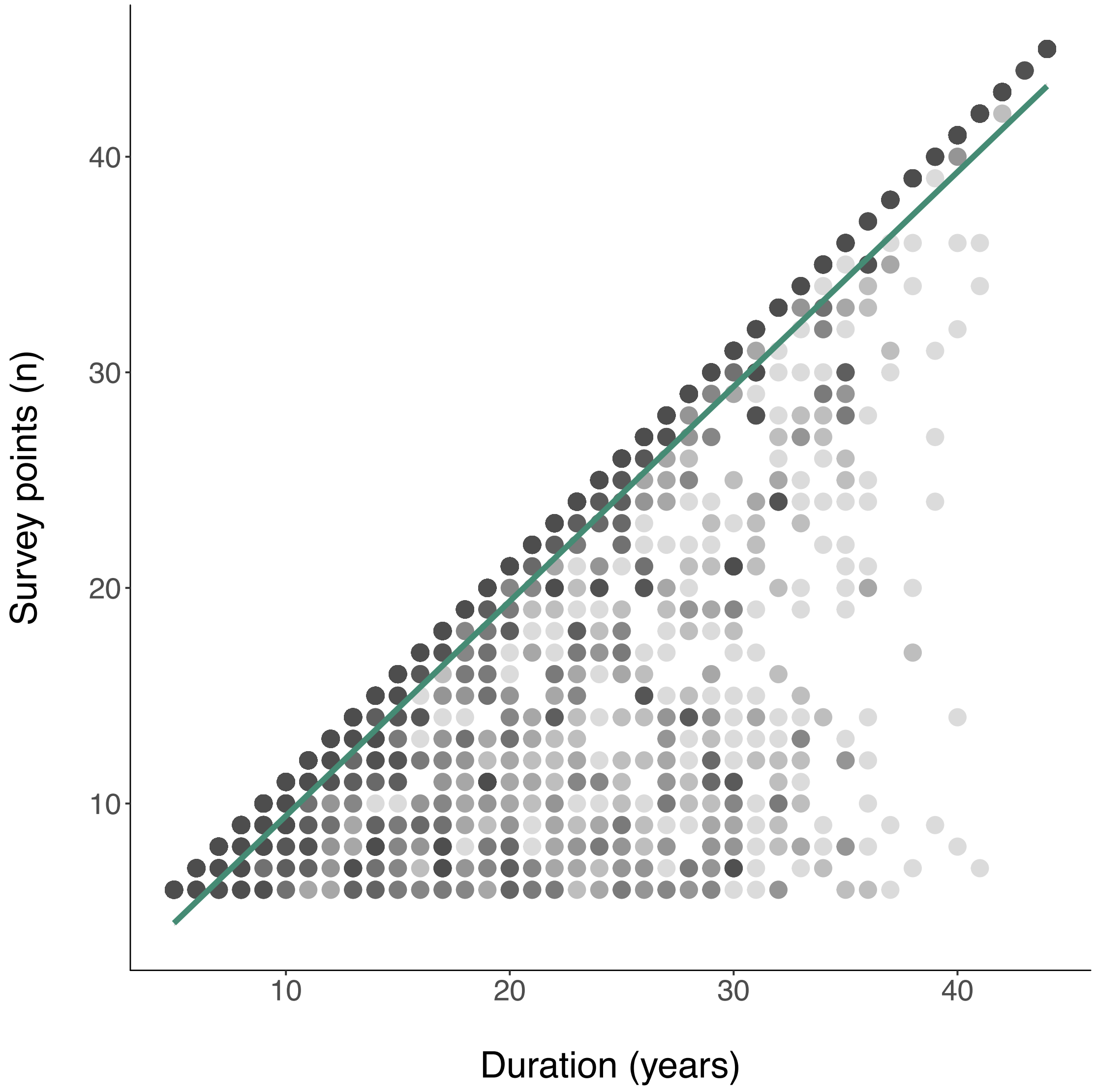


**Figure S8. Number of survey points within time-series positively correlates with time-series duration.** We included time-series with more than five survey points in time in our analyses, but populations were not always monitored in each intervening year. Green line shows a linear model fit of survey points versus duration. There was a minimum of six time points for each time-series. Among the time-series we analysed, 18% had a duration of less than 10 years, 30% had a duration between 10 and 20 years, 18% had a duration between 21 and 30 years, and 33% had a duration between 31 and 44 years. See Figure S3 for the density distribution of monitoring duration for the studies we included in our analyses.

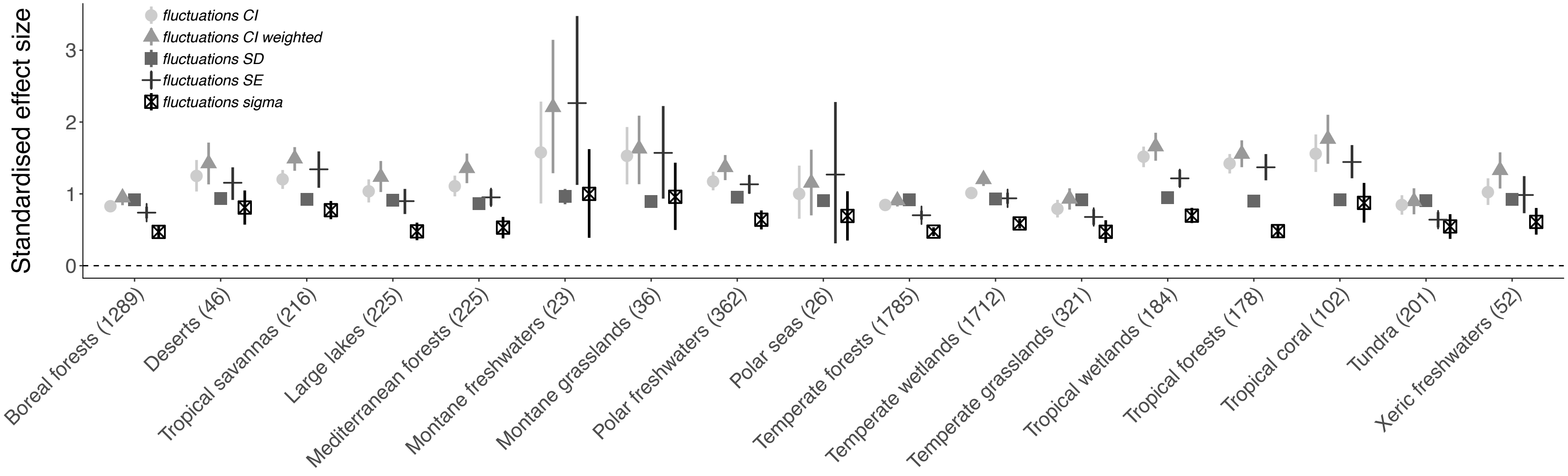


**Figure S9. Population fluctuations did not show distinct biome-specific patterns, with the exception of montane and tropical biomes where fluctuations were more pronounced compared to the rest of the biomes we studied.** The five estimates for each category refer to different analytical approaches, where the response variables in the models were: 1) the standard error around the slope estimates of the linear model fits of abundance versus year (circles), 2) half of the 95% confidence interval around the $\mu$ value of population change (triangles), 3) half of the 95% confidence interval around $\mu$ weighted by $\tau^2$, (full squares), 4) the process noise ($\sigma^2$) from the state-space models, and 5) the standard deviation of the raw data for each population time-series (empty squares). The process noise is the total variance around the population trend minus the variance attributed to observation error. The effect sizes were standardized by dividing the effect size by the standard deviation of the corresponding input data. Error bars show 95% credible intervals.

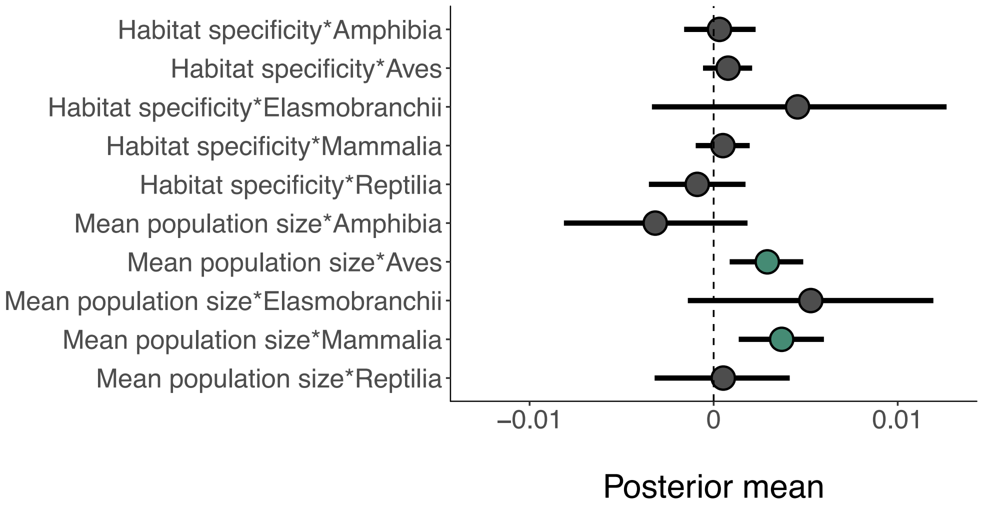


**Figure S10. Birds and mammals with larger mean population sizes are more likely to experience population increases.** We tested for interaction effects of rarity and taxa on population trends and, with the exception of mean population size for mammals and birds, rarity traits were not significant predictors of population change. Teal colour indicates posterior means where the 95% credible intervals did not overlap zero, grey colour indicates the opposite. Error bars show 95% credible intervals.

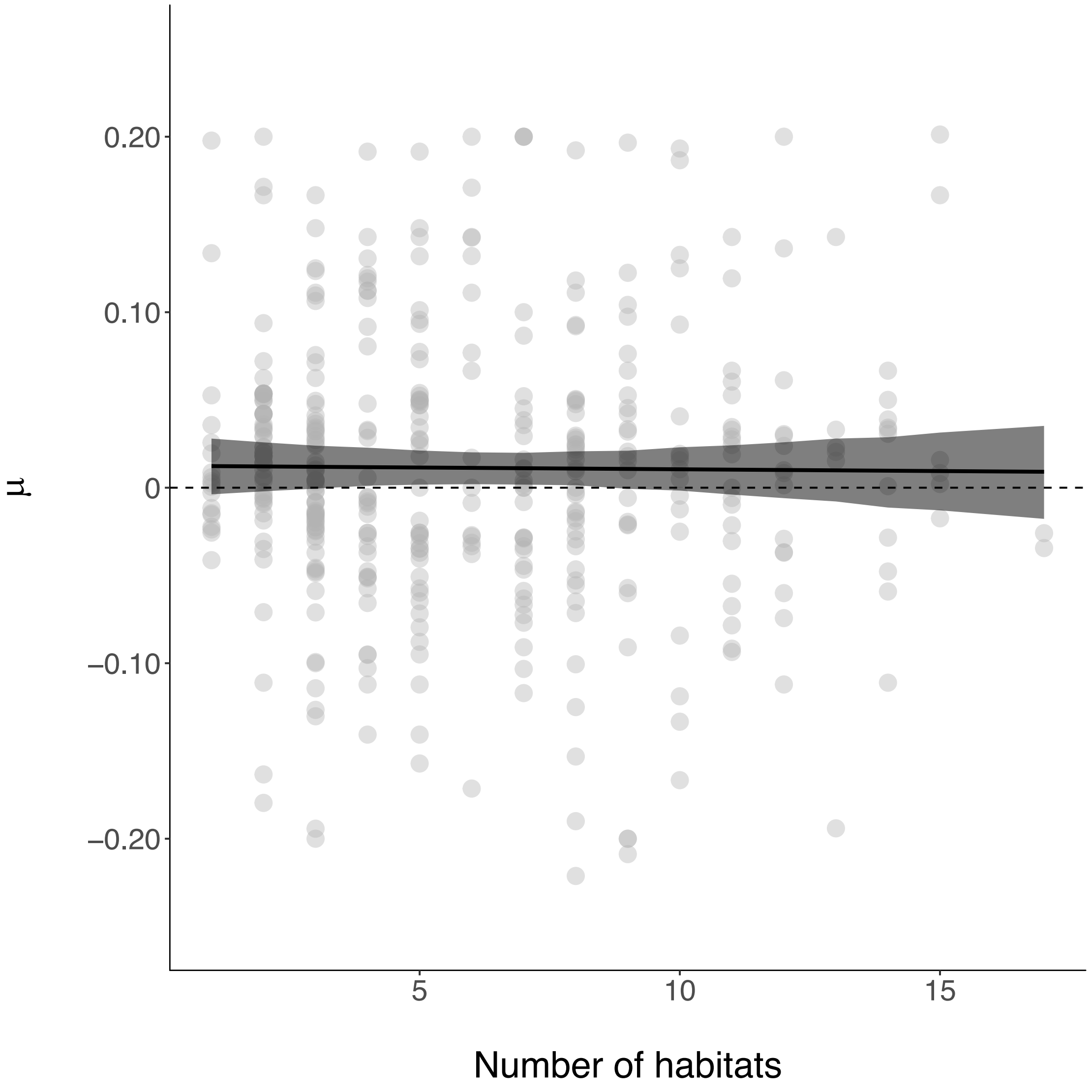


**Figure S11. Variation in vertebrate population trends was not explained by habitat specificity.** Habitat specificity was calculated as the number of different habitats occupied by each species which we derived by surveying the 'Habitat and Ecology' profile for each species on the IUCN Red List website. The $\mu$ values of population change are derived from state-space model fits of changes in abundance over the monitoring duration for each population. See Table S2 for full model outputs. This figure is based on populations monitored in the UK, see Figure 3c for the effects of habitat specificity on population trends across bird and mammal species globally.

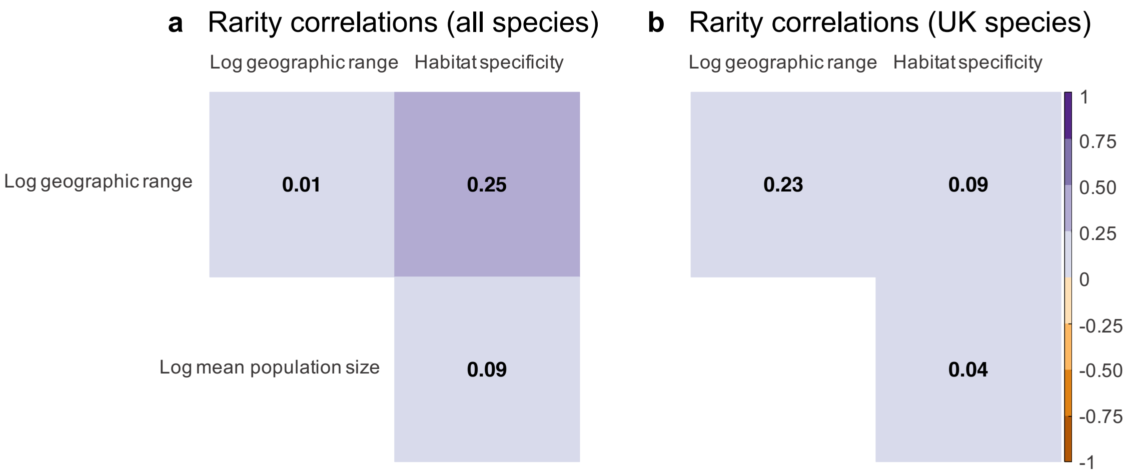


**Figure S12. The three rarity metrics used in this study were weakly correlated at both UK and global scales.** See Table S1 for sample sizes on each geographic scale.

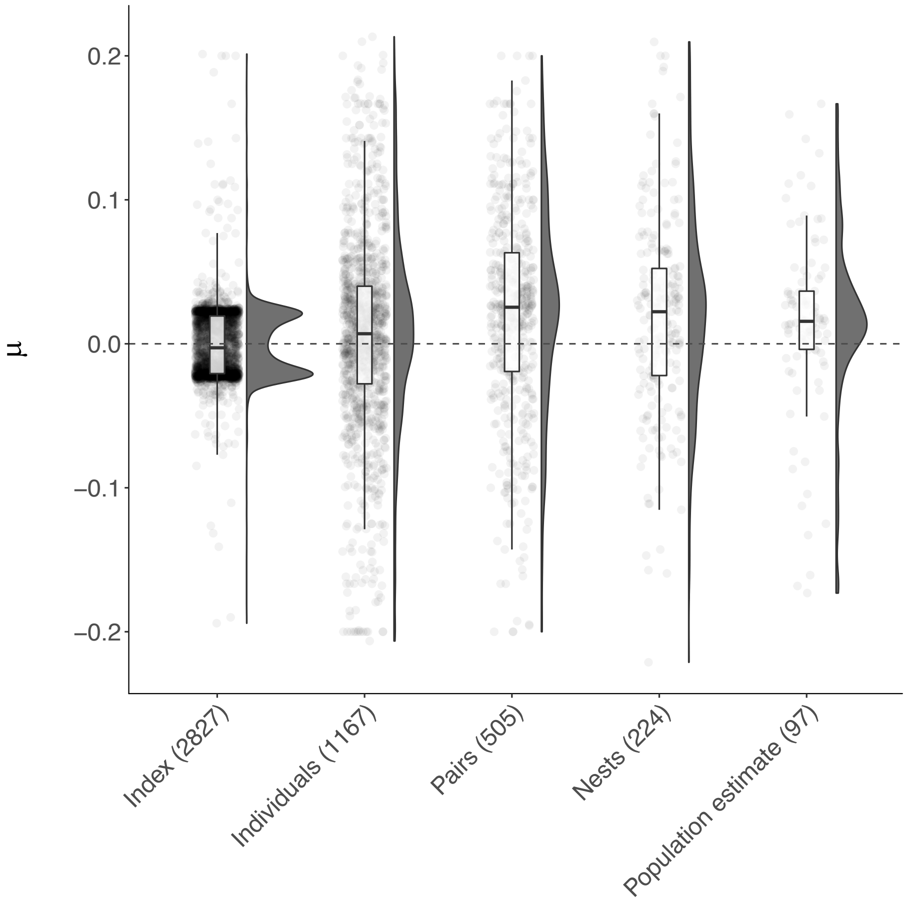


**Figure S13. There was no systematic difference in the distribution of population trends across commonly used abundance metrics in the Living Planet Database.** Population trends represent $\mu$ values from state-space Numbers on the x axis correspond to sample size in each category. Density plots show the density distribution of population trends across sampling units, points show the raw values and boxplots show the mean, first and third quartiles, with boxplot whiskers indicating the distance that covers 1.5 times the interquartile range. Population data where the units are an index were more likely to have weakly increasing or decreasing trends with many populations with $\mu$ values around 0.025 and -0.025. These $\mu$ estimates are reasonable trend estimates for these time series; however, there seem to be population within the Living Planet Database that are modelled, particularly for studies with longer duration which could partially explain these peaks in the population trend estimates near zero (see Figures 1, 2, S6 and S7).

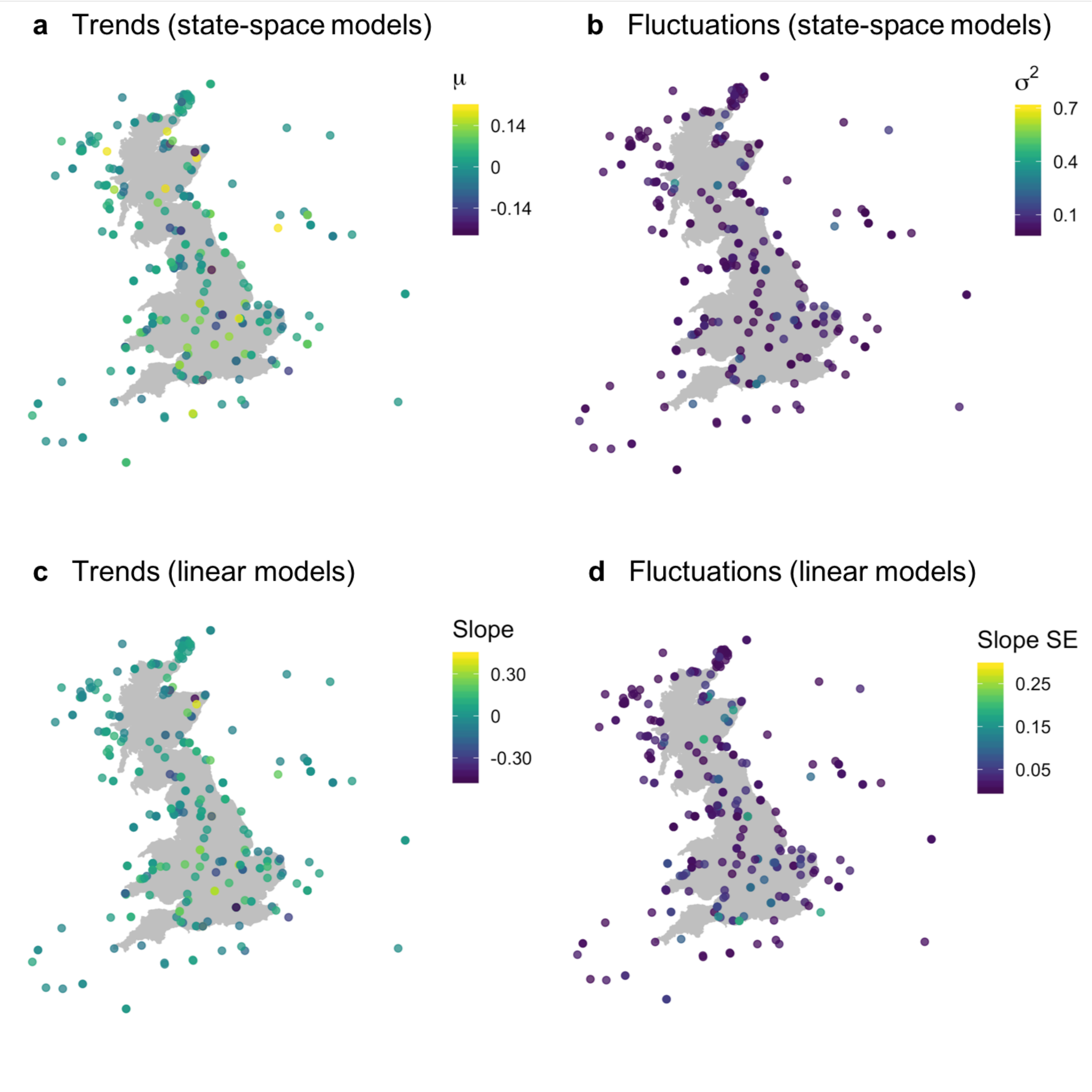


**Figure S14. Population trends and fluctuations did not show geographic patterning within the UK.** Results include 508 populations from 237 species in the UK. See methods for additional details on the different ways we quantified trends and fluctuations.

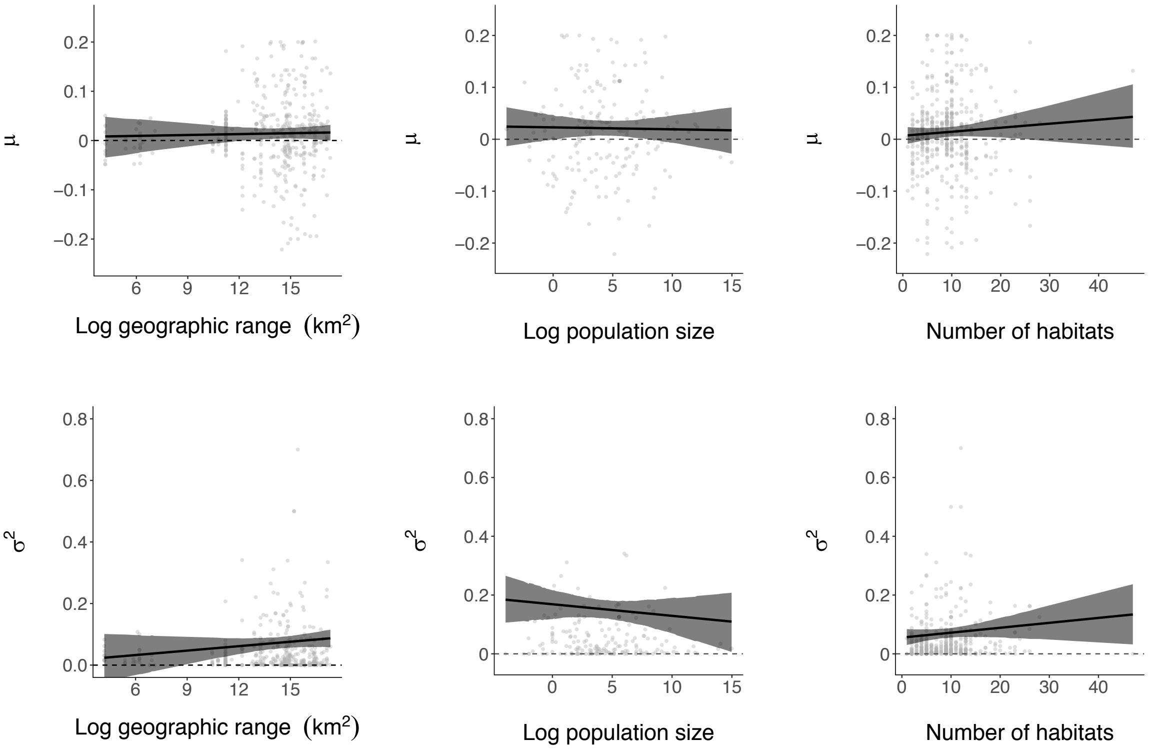


**Figure S15. The effects of rarity on population trends and fluctuations were consistent at both UK (pictured here) and global scales (Figure 3).** Rare species were not more likely to decline than common species. Populations from species with smaller mean population sizes and populations from habitat generalist species were more likely to fluctuate. The $\mu$ values of population trend (plots **a-c**) and the $\sigma^2$ values of population fluctuation (d-f) were derived from state-space model fits of changes in abundance over the monitoring duration for each population. The population fluctuations represent the process noise from the state-space models which is the total variance around the population trend minus the variance attributed to observation error. Lines on plots a-f show model fits and 95% credible intervals. See Table S1 for sample sizes for each analysis and Table S3 for model outputs.

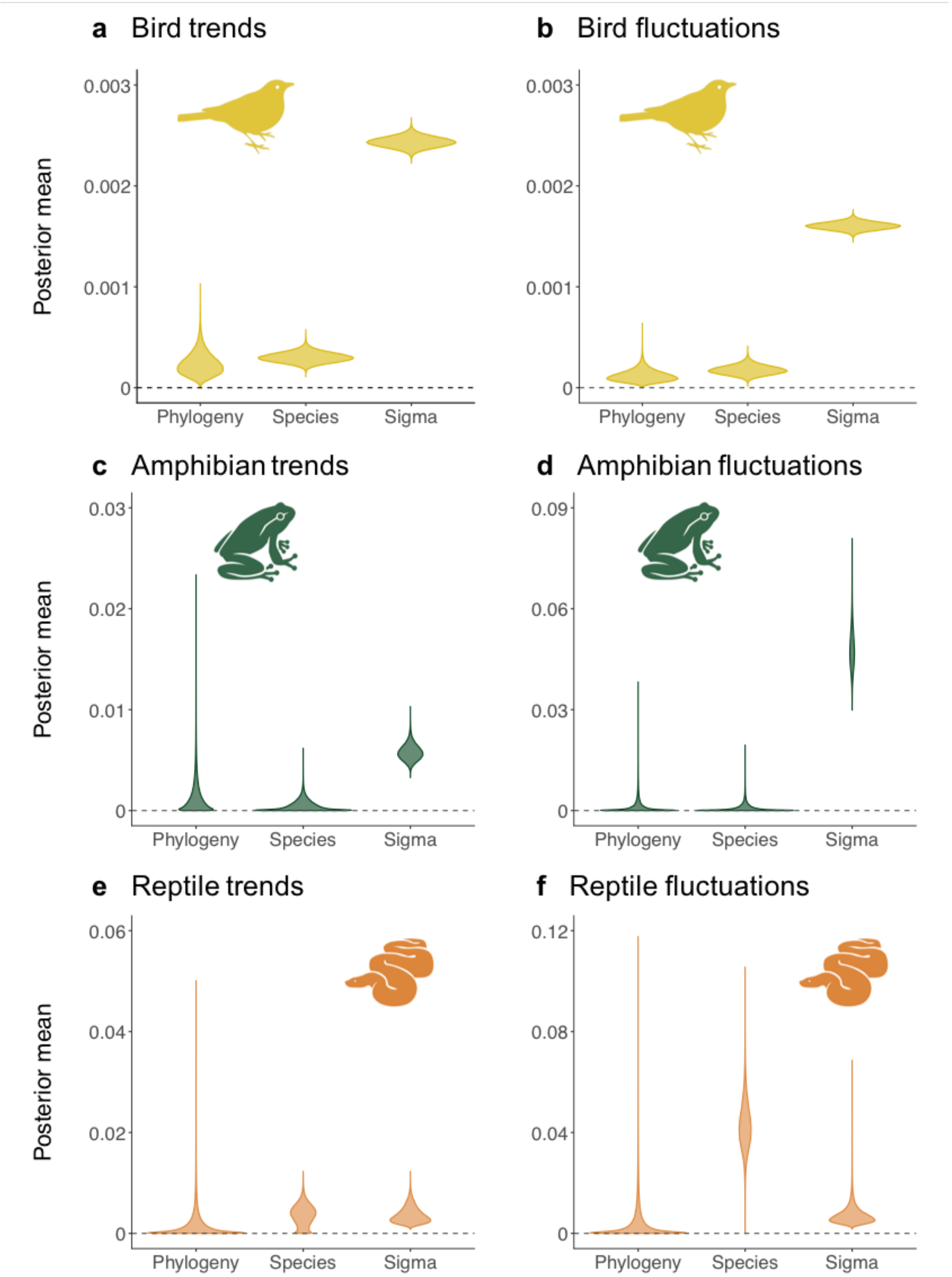


161

162 **Figure S16. There were no phylogenetic patterns in the population trends and**
163 **fluctuations of birds, amphibians and reptiles.** Phylogeny refers to variance explained by
164 phylogenetic relationships, species refers to variance explained by within-species differences
165 (some species were represented by more than one population, thus introducing species-level
166 variance), and residual variance refers to the variance not explained by phylogeny and species

effects. The figure shows violin plots of the distributions of posterior means for phylogenetic, species and residual variance across taxa (the wider the violin, the more records there are with that value). The distributions are based on ten models for each taxon for each type of population change using 10 random trees to account for phylogenetic uncertainty. Phylogeny effects were calculated based on a branch length covariance matrix. See Table S4 for full model outputs.

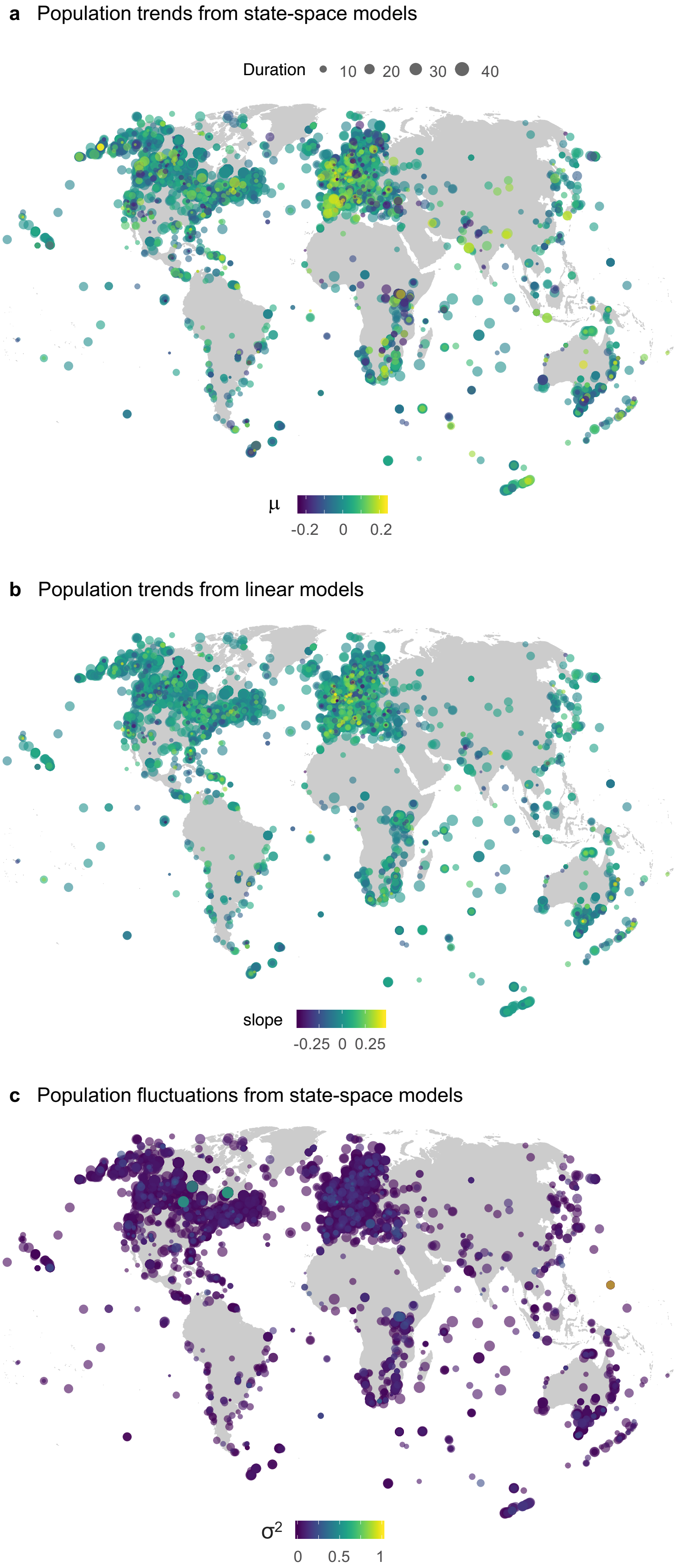
a Population trends from state-space models
Duration 10 20 30 40
μ
-0.2 0 0.2
b Population trends from linear models
slope
-0.25 0 0.25
c Population fluctuations from state-space models
σ2
0 0.5 1

**Figure S17. Population change is ubiquitous across the planet, with no distinct hotspots of declines, increases or fluctuations.** Maps show geographic distribution of time-series colour-coded by the magnitude of change experienced. The population fluctuations represent the process noise from the state-space models which is the total variance around the population trend minus the variance attributed to observation error. See methods for additional details on calculating trends and fluctuations.

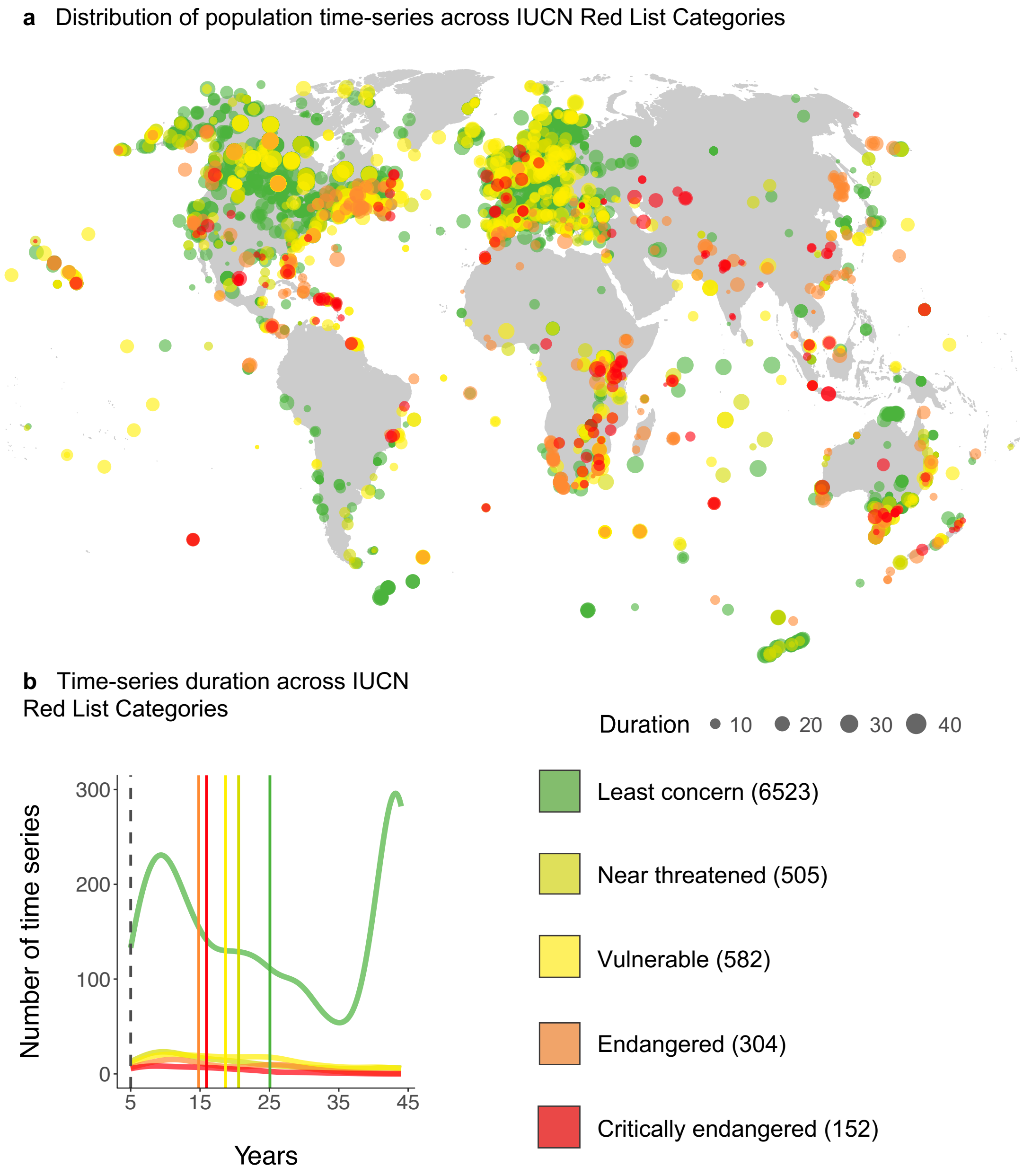


**Figure S18. Species across the whole spectrum of the IUCN Red List Categories are distributed around the world, with a concentration of least concern species in Northern America and Europe.** In our study, we included time-series with more than five survey points in time (b), with the dashed line five years and solid lines showing the mean duration for each category. Populations from least concern species were monitored for longer durations. Numbers next to each category show sample size.

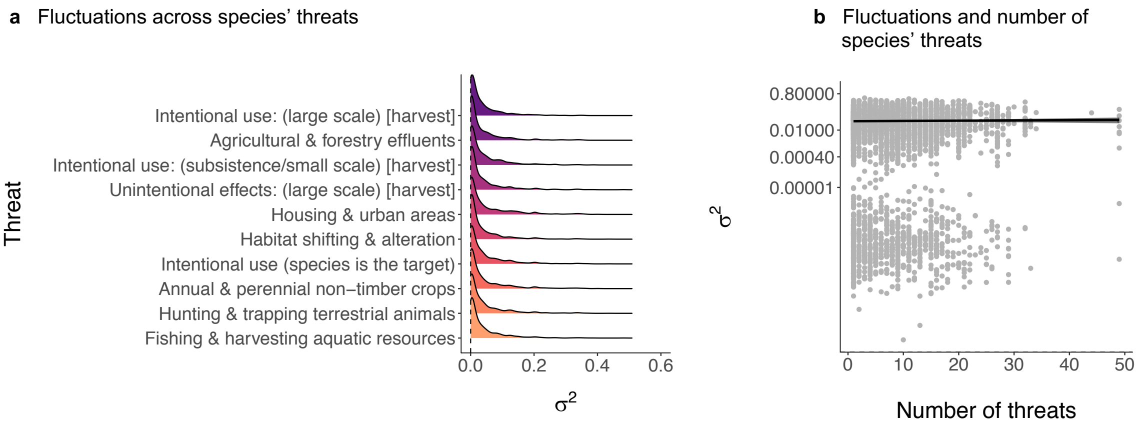


**Figure S19. Population fluctuations did not differ based on the type and number of threats based on species' IUCN Red List profiles.** Fluctuations were estimated using the process noise ($\sigma^2$) values from state-space model fits of changes in abundance over the monitoring duration for each population. Lines in (b) show model fit and 95% credible intervals, where "number of threats" refers to the number of different threats that each species, whose populations are locally monitored, could be exposed to throughout their distribution range, based on species' IUCN Red List profiles. See Methods for how the types of threats were derived and Table S2 for model outputs.

**Table S1. Number of species and populations included in analyses.**

| Scale | Analysis | Number of species | Number of populations | Number of species with >3 populations |
|---|---|---|---|---|
| **Global** | System | 2074 | 9286 | 834 |
| | Biome | 2074 | 9286 | 834 |
| | Taxa | | | |
| | - *Actinopterygii* | 544 | 1626 | 151 |
| | - *Amphibia* | 78 | 193 | 21 |
| | - *Aves* | 968 | 5852 | 538 |
| | - *Mammalia* | 306 | 1158 | 94 |
| | - *Reptilia* | 110 | 322 | 16 |
| | Red List status | 1702 | 8064 | 708 |
| | Latitude | 2074 | 9286 | 834 |
| | Duration | 2074 | 9286 | 834 |
| **UK** | Geographic range | 167 | 381 | 32 |
| | Population size | 112 | 253 | 19 |
| | Habitat specificity | 144 | 289 | 29 |
| | Red List status | 153 | 364 | 31 |

**Table S2. Model outputs from global analyses.** Sigma is the overall model residual variance. Net population change is estimated using $\mu$ values derived from state-space models of population abundance versus time and slopes of linear models of population abundance versus time. The weighted $\mu$ models included $\mu$ as a response variable, weighted by $\tau$, the observation error estimate derived from the state-space models. The fluctuation models included the process noise ($\sigma^2$) values from state-space models, half of the 95% confidence interval around the $\mu$ value of population change, the standard error around the slopes of linear models of population abundance versus time, and the standard deviation of the raw time-series data for each population. The process noise is a metric of population fluctuations, whereas the rest of the metrics show population variability. The weighted fluctuation models were weighted by $\tau$, the observation error estimate derived from the state-space models.

| Model name | Variable | Post. mean | Lower 95% CI | Upper 95% CI | Eff. sample size | pMCMC | Effect |
|---|---|---|---|---|---|---|---|
| **Latitude - $\mu$** | (Intercept) | 0.003 | 0.0002 | 0.005 | 18,000 | 0.037 | fixed |
| | Decimal.Latitude | 0.00001 | -0.00004 | 0.0001 | 18,000 | 0.687 | fixed |
| | sigma | 0.003 | 0.003 | 0.003 | 15,774 | | residual |
| **Realm - $\mu$** | Freshwater | 0.003 | -0.001 | 0.006 | 18,000 | 0.146 | fixed |
| | Marine | 0.004 | 0.0003 | 0.007 | 18,597 | 0.029 | fixed |
| | Terrestrial | 0.003 | 0.0002 | 0.005 | 18,000 | 0.035 | fixed |
| | sigma | 0.003 | 0.003 | 0.003 | 16,324 | | residual |
| **Realm - weighted** | Freshwater | 0.004 | -0.001 | 0.009 | 18,000 | 0.156 | fixed |
| | Marine | 0.003 | -0.002 | 0.008 | 18,000 | 0.168 | fixed |
| | Terrestrial | 0.003 | 0.00001 | 0.007 | 18,000 | 0.053 | fixed |
| | sigma | 0.003 | 0.003 | 0.003 | 13,851 | | residual |
| **Realm - slope** | Freshwater | 0.006 | -0.0002 | 0.012 | 18,000 | 0.067 | fixed |
| | Marine | 0.003 | -0.002 | 0.009 | 18,000 | 0.263 | fixed |
| | Terrestrial | 0.004 | -0.001 | 0.008 | 18,000 | 0.114 | fixed |
| | sigma | 0.008 | 0.008 | 0.008 | 15,950 | | residual |
| **Realm - fluctuations $\sigma$** | Terrestrial | 0.022 | 0.020 | 0.024 | 16,584 | 0.0001 | fixed |
| | Marine | 0.028 | 0.026 | 0.030 | 21,131 | 0.0001 | fixed |
| | Freshwater | 0.028 | 0.025 | 0.030 | 17,579 | 0.0001 | fixed |
| | sigma | 0.002 | 0.002 | 0.002 | 15,086 | | residual |

| | | | | | | | |
|---|---|---|---|---|---|---|---|
| **Realm - fluctuations CI** | Freshwater | 0.144 | 0.136 | 0.152 | 18,000 | 0.0001 | fixed |
| | Marine | 0.148 | 0.140 | 0.155 | 18,000 | 0.0001 | fixed |
| | Terrestrial | 0.118 | 0.111 | 0.124 | 18,000 | 0.0001 | fixed |
| | sigma | 0.012 | 0.011 | 0.012 | 17,555 | | residual |
| **Realm - fluctuations CI weighted** | Terrestrial | 0.130 | 0.122 | 0.138 | 18,000 | 0.0001 | fixed |
| | Marine | 0.169 | 0.159 | 0.180 | 18,000 | 0.0001 | fixed |
| | Freshwater | 0.170 | 0.159 | 0.181 | 16,896 | 0.0001 | fixed |
| | sigma | 0.011 | 0.010 | 0.011 | 18,000 | | residual |
| **Realm - fluctuations SE** | Freshwater | 0.135 | 0.126 | 0.143 | 18,000 | 0.0001 | fixed |
| | Marine | 0.139 | 0.131 | 0.148 | 18,000 | 0.0001 | fixed |
| | Terrestrial | 0.109 | 0.102 | 0.115 | 18,440 | 0.0001 | fixed |
| | sigma | 0.012 | 0.011 | 0.012 | 18,000 | | residual |
| **Realm - fluctuations SD** | Freshwater | 0.563 | 0.557 | 0.569 | 18,000 | 0.0001 | fixed |
| | Marine | 0.568 | 0.562 | 0.573 | 18,000 | 0.0001 | fixed |
| | Terrestrial | 0.547 | 0.542 | 0.552 | 18,000 | 0.0001 | fixed |
| | sigma | 0.008 | 0.008 | 0.008 | 16,850 | | residual |
| **Biome - $\mu$** | Boreal forests/taiga | 0.002 | -0.002 | 0.006 | 18,664 | 0.235 | fixed |
| | Deserts and xeric shrublands | -0.006 | -0.023 | 0.012 | 18,000 | 0.526 | fixed |
| | Trop. and subtrop. grasslands savannas and shrublands | -0.009 | -0.019 | 0.0004 | 18,000 | 0.061 | fixed |
| | Large lakes | 0.003 | -0.005 | 0.012 | 18,000 | 0.405 | fixed |
| | Mediterranean forests woodlands and scrub | 0.005 | -0.003 | 0.014 | 18,000 | 0.188 | fixed |
| | Montane freshwaters | 0.011 | -0.028 | 0.051 | 18,000 | 0.574 | fixed |
| | Montane grasslands and shrublands | 0.030 | 0.010 | 0.050 | 18,000 | 0.003 | fixed |
| | Polar freshwaters | 0.004 | -0.003 | 0.012 | 18,000 | 0.258 | fixed |
| | Polar seas | -0.011 | -0.035 | 0.014 | 18,000 | 0.397 | fixed |

| | | | | | | | |
|---|---|---|---|---|---|---|---|
| | Temperate forests | 0.002 | -0.002 | 0.005 | 18,000 | 0.318 | fixed |
| | Temperate wetlands and rivers | 0.002 | -0.002 | 0.007 | 18,000 | 0.226 | fixed |
| | Temperate grasslands savannas and shrublands | -0.004 | -0.011 | 0.003 | 18,000 | 0.247 | fixed |
| | Tropical wetlands and rivers | 0.002 | -0.007 | 0.011 | 18,000 | 0.725 | fixed |
| | Tropical and subtropical forests | 0.018 | 0.009 | 0.027 | 18,000 | 0.0001 | fixed |
| | Tropical coral | 0.024 | 0.012 | 0.037 | 18,000 | 0.0001 | fixed |
| | Tundra | 0.007 | -0.002 | 0.015 | 19,239 | 0.111 | fixed |
| | Xeric freshwaters and endorheic basins | -0.002 | -0.017 | 0.013 | 18,000 | 0.831 | fixed |
| | sigma | 0.003 | 0.003 | 0.003 | 15,790 | | residual |
| **Biome - slope** | Boreal forests/taiga | 0.002 | -0.004 | 0.007 | 18,000 | 0.523 | fixed |
| | Deserts and xeric shrublands | -0.027 | -0.050 | -0.002 | 18,000 | 0.031 | fixed |
| | Trop. and subtrop. grasslands savannas and shrublands | -0.017 | -0.030 | -0.004 | 18,000 | 0.014 | fixed |
| | Large lakes | 0.002 | -0.010 | 0.015 | 18,781 | 0.792 | fixed |
| | Mediterranean forests woodlands and scrub | 0.010 | -0.002 | 0.023 | 17,610 | 0.109 | fixed |
| | Montane freshwaters | 0.021 | -0.034 | 0.074 | 18,000 | 0.445 | fixed |
| | Montane grasslands and shrublands | 0.021 | -0.005 | 0.046 | 17,224 | 0.115 | fixed |
| | Polar freshwaters | 0.003 | -0.009 | 0.014 | 18,000 | 0.660 | fixed |
| | Polar seas | -0.016 | -0.046 | 0.015 | 17,196 | 0.320 | fixed |
| | Temperate forests | 0.003 | -0.001 | 0.008 | 17,408 | 0.165 | fixed |

| | | | | | | | |
|---|---|---|---|---|---|---|---|
| | Temperate wetlands and rivers | 0.004 | -0.001 | 0.010 | 18,000 | 0.123 | fixed |
| | Temperate grasslands savannas and shrublands | -0.001 | -0.010 | 0.007 | 18,000 | 0.780 | fixed |
| | Tropical wetlands and rivers | 0.003 | -0.011 | 0.016 | 18,000 | 0.713 | fixed |
| | Tropical and subtropical forests | 0.025 | 0.012 | 0.038 | 17,560 | 0.0002 | fixed |
| | Tropical coral | 0.036 | 0.019 | 0.052 | 18,000 | 0.0001 | fixed |
| | Tundra | 0.013 | -0.0003 | 0.026 | 17,563 | 0.048 | fixed |
| | Xeric freshwaters and endorheic basins | 0.014 | -0.010 | 0.038 | 18,257 | 0.265 | fixed |
| | sigma | 0.003 | 0.003 | 0.003 | 13,753 | | residual |
| **Biome - weighted** | Boreal forests/taiga | 0.003 | -0.003 | 0.010 | 18,000 | 0.334 | fixed |
| | Deserts and xeric shrublands | -0.019 | -0.049 | 0.011 | 18,000 | 0.208 | fixed |
| | Trop. and subtrop. grasslands savannas and shrublands | -0.004 | -0.021 | 0.013 | 18,000 | 0.635 | fixed |
| | Large lakes | 0.007 | -0.007 | 0.021 | 18,000 | 0.336 | fixed |
| | Mediterranean forests woodlands and scrub | 0.006 | -0.008 | 0.020 | 19,170 | 0.402 | fixed |
| | Montane freshwaters | 0.023 | -0.044 | 0.094 | 18,000 | 0.521 | fixed |
| | Montane grasslands and shrublands | 0.037 | 0.003 | 0.071 | 20,405 | 0.033 | fixed |
| | Polar freshwaters | 0.012 | -0.001 | 0.025 | 18,000 | 0.069 | fixed |
| | Polar seas | -0.023 | -0.066 | 0.018 | 18,000 | 0.292 | fixed |
| | Temperate forests | 0.003 | -0.003 | 0.009 | 18,000 | 0.265 | fixed |

| | | | | | | | |
|---|---|---|---|---|---|---|---|
| | Temperate wetlands and rivers | 0.005 | -0.002 | 0.012 | 17,585 | 0.169 | fixed |
| | Temperate grasslands savannas and shrublands | -0.008 | -0.019 | 0.004 | 18,000 | 0.161 | fixed |
| | Tropical wetlands and rivers | 0.006 | -0.010 | 0.022 | 18,000 | 0.473 | fixed |
| | Tropical and subtropical forests | 0.014 | -0.001 | 0.029 | 18,000 | 0.068 | fixed |
| | Tropical coral | 0.039 | 0.018 | 0.060 | 18,000 | 0.0003 | fixed |
| | Tundra | 0.013 | -0.001 | 0.027 | 18,000 | 0.071 | fixed |
| | Xeric freshwaters and endorheic basins | -0.021 | -0.048 | 0.003 | 18,000 | 0.099 | fixed |
| | sigma | 0.008 | 0.007 | 0.008 | 16,533 | | residual |
| **Biome - fluctuations *σ*** | Boreal forests/taiga | 0.018 | 0.015 | 0.021 | 18,000 | 0.0001 | fixed |
| | Deserts and xeric shrublands | 0.044 | 0.031 | 0.057 | 18,000 | 0.0001 | fixed |
| | Trop. and subtrop. grasslands savannas and shrublands | 0.044 | 0.037 | 0.051 | 18,000 | 0.0001 | fixed |
| | Large lakes | 0.024 | 0.018 | 0.030 | 18,000 | 0.0001 | fixed |
| | Mediterranean forests woodlands and scrub | 0.022 | 0.016 | 0.028 | 18,000 | 0.0001 | fixed |
| | Montane freshwaters | 0.047 | 0.018 | 0.075 | 18,000 | 0.001 | fixed |
| | Montane grasslands and shrublands | 0.031 | 0.016 | 0.046 | 18,000 | 0.0001 | fixed |
| | Polar freshwaters | 0.027 | 0.021 | 0.033 | 17,652 | 0.0001 | fixed |
| | Polar seas | 0.037 | 0.019 | 0.055 | 17,689 | 0.0001 | fixed |
| | Temperate forests | 0.019 | 0.017 | 0.022 | 17,789 | 0.0001 | fixed |

| | | | | | | |
|---|---|---|---|---|---|---|
| | Temperate wetlands and rivers | 0.024 | 0.022 | 0.027 | 18,000 | 0.0001 | fixed |
| | Temperate grasslands savannas and shrublands | 0.015 | 0.010 | 0.020 | 18,000 | 0.0001 | fixed |
| | Tropical wetlands and rivers | 0.046 | 0.039 | 0.054 | 18,000 | 0.0001 | fixed |
| | Tropical and subtropical forests | 0.041 | 0.035 | 0.048 | 17,508 | 0.0001 | fixed |
| | Tropical coral | 0.029 | 0.020 | 0.038 | 18,000 | 0.0001 | fixed |
| | Tundra | 0.020 | 0.014 | 0.026 | 18,497 | 0.0001 | fixed |
| | Xeric freshwaters and endorheic basins | 0.039 | 0.028 | 0.052 | 18,845 | 0.0001 | fixed |
| | sigma | 0.002 | 0.002 | 0.002 | 14,440 | | residual |
| **Biome - fluctuations CI** | Boreal forests/taiga | 0.091 | 0.083 | 0.100 | 18,000 | 0.0001 | fixed |
| | Deserts and xeric shrublands | 0.210 | 0.173 | 0.247 | 17,389 | 0.0001 | fixed |
| | Trop. and subtrop. grasslands savannas and shrublands | 0.200 | 0.178 | 0.222 | 18,000 | 0.0001 | fixed |
| | Large lakes | 0.112 | 0.095 | 0.130 | 18,000 | 0.0001 | fixed |
| | Mediterranean forests woodlands and scrub | 0.132 | 0.115 | 0.150 | 18,000 | 0.0001 | fixed |
| | Montane freshwaters | 0.197 | 0.108 | 0.286 | 18,000 | 0.0001 | fixed |
| | Montane grasslands and shrublands | 0.163 | 0.121 | 0.206 | 18,000 | 0.0001 | fixed |
| | Polar freshwaters | 0.147 | 0.132 | 0.164 | 18,000 | 0.0001 | fixed |
| | Polar seas | 0.146 | 0.096 | 0.204 | 17,894 | 0.0001 | fixed |
| | Temperate forests | 0.099 | 0.091 | 0.107 | 18,000 | 0.0001 | fixed |

| | | | | | | | |
|---|---|---|---|---|---|---|---|
| | Temperate wetlands and rivers | 0.132 | 0.123 | 0.141 | 18,000 | 0.0001 | fixed |
| | Temperate grasslands savannas and shrublands | 0.092 | 0.077 | 0.106 | 18,000 | 0.0001 | fixed |
| | Tropical wetlands and rivers | 0.219 | 0.198 | 0.239 | 18,000 | 0.0001 | fixed |
| | Tropical and subtropical forests | 0.197 | 0.178 | 0.215 | 18,405 | 0.0001 | fixed |
| | Tropical coral | 0.161 | 0.135 | 0.189 | 18,000 | 0.0001 | fixed |
| | Tundra | 0.107 | 0.090 | 0.124 | 18,338 | 0.0001 | fixed |
| | Xeric freshwaters and endorheic basins | 0.177 | 0.146 | 0.211 | 18,000 | 0.0001 | fixed |
| | sigma | 0.011 | 0.011 | 0.011 | 17,482 | | residual |
| **Biome - fluctuations CI weighted** | Boreal forests/taiga | 0.104 | 0.092 | 0.115 | 18,000 | 0.0001 | fixed |
| | Deserts and xeric shrublands | 0.239 | 0.190 | 0.288 | 18,524 | 0.0001 | fixed |
| | Trop. and subtrop. grasslands savannas and shrublands | 0.248 | 0.220 | 0.274 | 18,431 | 0.0001 | fixed |
| | Large lakes | 0.134 | 0.111 | 0.158 | 16,663 | 0.0001 | fixed |
| | Mediterranean forests woodlands and scrub | 0.162 | 0.137 | 0.187 | 18,000 | 0.0001 | fixed |
| | Montane freshwaters | 0.276 | 0.161 | 0.393 | 18,000 | 0.0001 | fixed |
| | Montane grasslands and shrublands | 0.173 | 0.121 | 0.222 | 17,341 | 0.0001 | fixed |
| | Polar freshwaters | 0.172 | 0.149 | 0.193 | 18,386 | 0.0001 | fixed |
| | Polar seas | 0.169 | 0.103 | 0.237 | 18,368 | 0.0001 | fixed |
| | Temperate forests | 0.107 | 0.097 | 0.116 | 18,000 | 0.0001 | fixed |

| | | | | | | | |
|---|---|---|---|---|---|---|---|
| | Temperate wetlands and rivers | 0.156 | 0.144 | 0.169 | 18,000 | 0.0001 | fixed |
| | Temperate grasslands savannas and shrublands | 0.107 | 0.090 | 0.124 | 18,000 | 0.0001 | fixed |
| | Tropical wetlands and rivers | 0.239 | 0.211 | 0.267 | 18,000 | 0.0001 | fixed |
| | Tropical and subtropical forests | 0.216 | 0.190 | 0.242 | 18,000 | 0.0001 | fixed |
| | Tropical coral | 0.183 | 0.147 | 0.218 | 18,000 | 0.0001 | fixed |
| | Tundra | 0.114 | 0.091 | 0.137 | 18,000 | 0.0001 | fixed |
| | Xeric freshwaters and endorheic basins | 0.230 | 0.186 | 0.273 | 18,000 | 0.0001 | fixed |
| | sigma | 0.010 | 0.010 | 0.011 | 16,732 | | residual |
| **Biome - fluctuations SE** | Boreal forests/taiga | 0.086 | 0.077 | 0.096 | 18,000 | 0.0001 | fixed |
| | Deserts and xeric shrublands | 0.204 | 0.162 | 0.242 | 18,000 | 0.0001 | fixed |
| | Trop. and subtrop. grasslands savannas and shrublands | 0.126 | 0.102 | 0.150 | 18,000 | 0.0001 | fixed |
| | Large lakes | 0.095 | 0.076 | 0.114 | 17,101 | 0.0001 | fixed |
| | Mediterranean forests woodlands and scrub | 0.149 | 0.131 | 0.167 | 17,283 | 0.0001 | fixed |
| | Montane freshwaters | 0.184 | 0.091 | 0.283 | 18,000 | 0.0002 | fixed |
| | Montane grasslands and shrublands | 0.110 | 0.065 | 0.156 | 18,000 | 0.0001 | fixed |
| | Polar freshwaters | 0.152 | 0.134 | 0.169 | 18,000 | 0.0001 | fixed |
| | Polar seas | 0.076 | 0.019 | 0.137 | 18,000 | 0.011 | fixed |
| | Temperate forests | 0.094 | 0.086 | 0.103 | 18,000 | 0.0001 | fixed |

| | | | | | | | |
|---|---|---|---|---|---|---|---|
| | Temperate wetlands and rivers | 0.123 | 0.113 | 0.133 | 18,000 | 0.0001 | fixed |
| | Temperate grasslands savannas and shrublands | 0.096 | 0.080 | 0.111 | 18,000 | 0.0001 | fixed |
| | Tropical wetlands and rivers | 0.229 | 0.207 | 0.251 | 18,000 | 0.0001 | fixed |
| | Tropical and subtropical forests | 0.154 | 0.133 | 0.174 | 18,000 | 0.0001 | fixed |
| | Tropical coral | 0.187 | 0.158 | 0.218 | 18,580 | 0.0001 | fixed |
| | Tundra | 0.111 | 0.092 | 0.129 | 18,663 | 0.0001 | fixed |
| | Xeric freshwaters and endorheic basins | 0.129 | 0.095 | 0.163 | 18,000 | 0.0001 | fixed |
| | sigma | 0.012 | 0.011 | 0.012 | 17,110 | | residual |
| **Biome - fluctuations SD** | Boreal forests/taiga | 0.523 | 0.517 | 0.530 | 18,000 | 0.0001 | fixed |
| | Deserts and xeric shrublands | 0.585 | 0.556 | 0.614 | 18,000 | 0.0001 | fixed |
| | Trop. and subtrop. grasslands savannas and shrublands | 0.602 | 0.585 | 0.619 | 18,000 | 0.0001 | fixed |
| | Large lakes | 0.540 | 0.527 | 0.555 | 18,000 | 0.0001 | fixed |
| | Mediterranean forests woodlands and scrub | 0.574 | 0.560 | 0.587 | 18,300 | 0.0001 | fixed |
| | Montane freshwaters | 0.605 | 0.537 | 0.671 | 18,000 | 0.0001 | fixed |
| | Montane grasslands and shrublands | 0.583 | 0.549 | 0.616 | 18,000 | 0.0001 | fixed |
| | Polar freshwaters | 0.570 | 0.557 | 0.582 | 18,000 | 0.0001 | fixed |
| | Polar seas | 0.548 | 0.504 | 0.588 | 18,000 | 0.0001 | fixed |
| | Temperate forests | 0.536 | 0.531 | 0.542 | 18,000 | 0.0001 | fixed |

| | | | | | | | |
|---|---|---|---|---|---|---|---|
| | Temperate wetlands and rivers | 0.557 | 0.551 | 0.564 | 18,000 | 0.0001 | fixed |
| | Temperate grasslands savannas and shrublands | 0.526 | 0.515 | 0.537 | 18,000 | 0.0001 | fixed |
| | Tropical wetlands and rivers | 0.608 | 0.593 | 0.625 | 18,000 | 0.0001 | fixed |
| | Tropical and subtropical forests | 0.614 | 0.600 | 0.629 | 18,000 | 0.0001 | fixed |
| | Tropical coral | 0.609 | 0.588 | 0.629 | 18,264 | 0.0001 | fixed |
| | Tundra | 0.539 | 0.525 | 0.553 | 18,000 | 0.0001 | fixed |
| | Xeric freshwaters and endorheic basins | 0.556 | 0.530 | 0.580 | 18,000 | 0.0001 | fixed |
| | sigma | 0.007 | 0.007 | 0.008 | 17,425 | | residual |
| **Taxa - *μ*** | Actinopterygii | 0.00004 | -0.004 | 0.004 | 18,851 | 0.986 | fixed |
| | Amphibia | -0.012 | -0.022 | -0.002 | 17,369 | 0.027 | fixed |
| | Aves | 0.003 | 0.001 | 0.006 | 18,000 | 0.007 | fixed |
| | Elasmobranchii | -0.010 | -0.022 | 0.002 | 17,095 | 0.097 | fixed |
| | Mammalia | 0.010 | 0.005 | 0.015 | 19,689 | 0.0001 | fixed |
| | Reptilia | 0.010 | 0.001 | 0.020 | 18,000 | 0.029 | fixed |
| | sigma | 0.003 | 0.003 | 0.003 | 16,346 | | residual |
| **Taxa - weighted** | Actinopterygii | -0.001 | -0.007 | 0.004 | 18,000 | 0.744 | fixed |
| | Amphibia | -0.016 | -0.032 | -0.001 | 18,000 | 0.046 | fixed |
| | Aves | 0.005 | 0.002 | 0.008 | 18,000 | 0.003 | fixed |
| | Elasmobranchii | -0.017 | -0.035 | 0.002 | 18,000 | 0.077 | fixed |
| | Mammalia | 0.011 | 0.004 | 0.017 | 18,490 | 0.001 | fixed |
| | Reptilia | 0.004 | -0.010 | 0.018 | 18,000 | 0.578 | fixed |
| | sigma | 0.003 | 0.003 | 0.003 | 14,129 | | residual |
| **Taxa - slope** | Actinopterygii | -0.001 | -0.008 | 0.005 | 18,000 | 0.664 | fixed |
| | Amphibia | -0.020 | -0.037 | -0.002 | 16,874 | 0.026 | fixed |
| | Aves | 0.006 | 0.001 | 0.010 | 18,000 | 0.009 | fixed |
| | Elasmobranchii | -0.018 | -0.039 | 0.002 | 17,613 | 0.083 | fixed |
| | Mammalia | 0.011 | 0.003 | 0.020 | 18,000 | 0.012 | fixed |
| | Reptilia | 0.022 | 0.006 | 0.038 | 18,000 | 0.007 | fixed |
| | sigma | 0.008 | 0.008 | 0.008 | 16,177 | | residual |

| | | | | | | | |
|---|---|---|---|---|---|---|---|
| **Taxa - fluctuations *σ*** | Actinopterygii | 0.032 | 0.030 | 0.035 | 17,489 | 0.0001 | fixed |
| | Amphibia | 0.040 | 0.033 | 0.047 | 18,000 | 0.0001 | fixed |
| | Aves | 0.018 | 0.017 | 0.020 | 18,000 | 0.0001 | fixed |
| | Elasmobranchii | 0.030 | 0.022 | 0.039 | 18,000 | 0.0001 | fixed |
| | Mammalia | 0.035 | 0.032 | 0.038 | 17,868 | 0.0001 | fixed |
| | Reptilia | 0.034 | 0.028 | 0.041 | 18,000 | 0.0001 | fixed |
| | sigma | 0.002 | 0.002 | 0.002 | 13,289 | | residual |
| **Taxa - fluctuations CI** | Actinopterygii | 0.163 | 0.155 | 0.171 | 18,000 | 0.0001 | fixed |
| | Amphibia | 0.208 | 0.186 | 0.231 | 18,000 | 0.0001 | fixed |
| | Aves | 0.094 | 0.089 | 0.099 | 17,127 | 0.0001 | fixed |
| | Elasmobranchii | 0.152 | 0.125 | 0.177 | 18,000 | 0.0001 | fixed |
| | Mammalia | 0.182 | 0.172 | 0.193 | 18,000 | 0.0001 | fixed |
| | Reptilia | 0.195 | 0.176 | 0.216 | 18,476 | 0.0001 | fixed |
| | sigma | 0.012 | 0.011 | 0.012 | 18,000 | | residual |
| **Taxa - fluctuations CI weighted** | Actinopterygii | 0.191 | 0.180 | 0.202 | 18,000 | 0.0001 | fixed |
| | Amphibia | 0.272 | 0.242 | 0.304 | 17,535 | 0.0001 | fixed |
| | Aves | 0.105 | 0.098 | 0.112 | 18,000 | 0.0001 | fixed |
| | Elasmobranchii | 0.197 | 0.160 | 0.232 | 18,434 | 0.0001 | fixed |
| | Mammalia | 0.212 | 0.198 | 0.226 | 19,200 | 0.0001 | fixed |
| | Reptilia | 0.229 | 0.201 | 0.257 | 18,000 | 0.0001 | fixed |
| | sigma | 0.011 | 0.011 | 0.012 | 17,175 | | residual |
| **Taxa - fluctuations SE** | Actinopterygii | 0.159 | 0.150 | 0.168 | 18,000 | 0.0001 | fixed |
| | Amphibia | 0.224 | 0.199 | 0.247 | 17,493 | 0.0001 | fixed |
| | Aves | 0.080 | 0.074 | 0.086 | 18,000 | 0.0001 | fixed |
| | Elasmobranchii | 0.151 | 0.124 | 0.179 | 18,000 | 0.0001 | fixed |
| | Mammalia | 0.157 | 0.145 | 0.169 | 18,000 | 0.0001 | fixed |
| | Reptilia | 0.233 | 0.211 | 0.255 | 18,000 | 0.0001 | fixed |
| | sigma | 0.011 | 0.011 | 0.012 | 18,000 | | residual |
| **Taxa - fluctuations SD** | Actinopterygii | 0.572 | 0.566 | 0.579 | 18,000 | 0.0001 | fixed |
| | Amphibia | 0.612 | 0.596 | 0.630 | 18,000 | 0.0001 | fixed |
| | Aves | 0.530 | 0.526 | 0.534 | 18,000 | 0.0001 | fixed |
| | Elasmobranchii | 0.563 | 0.542 | 0.582 | 18,000 | 0.0001 | fixed |
| | Mammalia | 0.603 | 0.595 | 0.611 | 18,000 | 0.0001 | fixed |
| | Reptilia | 0.610 | 0.595 | 0.625 | 18,000 | 0.0001 | fixed |
| | sigma | 0.008 | 0.007 | 0.008 | 18,000 | | residual |

| | | | | | | | |
|---|---|---|---|---|---|---|---|
| **Number of time points - *μ*** | points | 0.00002 | -0.00004 | 0.0001 | 18,000 | 0.464 | fixed |
| | sigma | 0.003 | 0.003 | 0.003 | 16,697 | | residual |
| **Number of time points – *σ* (fluctuations)** | points | 0.0002 | 0.0001 | 0.0003 | 16,224 | 0.0001 | fixed |
| | sigma | 0.002 | 0.002 | 0.002 | 16,816 | | residual |
| **Duration** | duration | 0.0001 | 0.00003 | 0.0001 | 18,398 | 0.001 | fixed |
| | sigma | 0.003 | 0.003 | 0.004 | 18,000 | | residual |
| **Duration * System interaction** | Freshwater | 0.006 | 0.002 | 0.010 | 17,386 | 0.008 | fixed |
| | Marine | 0.009 | 0.004 | 0.014 | 18,000 | 0.001 | fixed |
| | Terrestrial | 0.009 | 0.006 | 0.013 | 18,000 | 0.0001 | fixed |
| | duration | -0.0001 | -0.0002 | 0.0001 | 18,025 | 0.532 | fixed |
| | Marine:duration | -0.0002 | -0.001 | 0.0001 | 18,000 | 0.162 | fixed |
| | Terrestrial:duration | -0.0002 | -0.0004 | 0.00004 | 17,941 | 0.113 | fixed |
| | sigma | 0.003 | 0.003 | 0.004 | 18,062 | | residual |
| **Duration * Taxa interaction** | Actinopterygii | 0.0005 | -0.006 | 0.007 | 18,000 | 0.890 | fixed |
| | Amphibia | -0.019 | -0.039 | 0.002 | 18,000 | 0.077 | fixed |
| | Aves | 0.010 | 0.006 | 0.014 | 18,000 | 0.0001 | fixed |
| | Mammalia | 0.011 | 0.003 | 0.019 | 18,000 | 0.009 | fixed |
| | Reptilia | 0.023 | 0.010 | 0.038 | 18,000 | 0.001 | fixed |
| | duration | -0.00003 | -0.0004 | 0.0003 | 18,000 | 0.879 | fixed |
| | Amphibia:duration | 0.001 | -0.001 | 0.002 | 18,000 | 0.444 | fixed |
| | Aves:duration | -0.0002 | -0.001 | 0.0001 | 18,000 | 0.242 | fixed |
| | Mammalia:duration | -0.00001 | -0.001 | 0.001 | 18,000 | 0.956 | fixed |
| | Reptilia:duration | -0.001 | -0.002 | -0.0001 | 18,000 | 0.026 | fixed |
| | sigma | 0.003 | 0.003 | 0.003 | 16,297 | | residual |
| **Sampling units - *μ*** | (Intercept) | -0.0001 | -0.003 | 0.003 | 18,000 | 0.918 | fixed |
| | Individuals (1167) | 0.006 | 0.002 | 0.010 | 18,000 | 0.003 | fixed |
| | Pairs (505) | 0.015 | 0.009 | 0.020 | 18,000 | 0.0001 | fixed |
| | Nests (224) | 0.015 | 0.007 | 0.023 | 18,000 | 0.0002 | fixed |
| | Population estimate (97) | 0.017 | 0.007 | 0.027 | 18,000 | 0.001 | fixed |
| | sigma | 0.002 | 0.002 | 0.002 | 15,983 | | residual |

| | | | | | | | |
|---|---|---|---|---|---|---|---|
| **Geographic range birds/mammals - *μ*** | (Intercept) | 0.012 | -0.002 | 0.026 | 18,000 | 0.082 | fixed |
| | log(Geographic range) | -0.0005 | -0.001 | 0.0004 | 18,000 | 0.278 | fixed |
| | sigma | 0.003 | 0.003 | 0.003 | 17,575 | | residual |
| **Geographic range birds/mammals - *μ* * taxa interaction** | (Intercept) | 0.068 | 0.031 | 0.107 | 18,000 | 0.0002 | fixed |
| | log(Geographic range) | -0.004 | -0.007 | -0.002 | 18,000 | 0.001 | fixed |
| | Aves | -0.076 | -0.119 | -0.031 | 18,000 | 0.001 | fixed |
| | log(Geographic range):Aves | 0.005 | 0.002 | 0.008 | 18,000 | 0.001 | fixed |
| | sigma | 0.003 | 0.003 | 0.003 | 16,888 | | residual |
| **Geographic range birds/mammals - weighted** | (Intercept) | 0.013 | -0.011 | 0.038 | 18,000 | 0.295 | fixed |
| | log(Geographic range) | -0.0004 | -0.002 | 0.001 | 18,000 | 0.564 | fixed |
| | sigma | 0.003 | 0.003 | 0.003 | 16,791 | | residual |
| **Geographic range birds/mammals - slope** | (Intercept) | 0.010 | -0.020 | 0.041 | 18,433 | 0.515 | fixed |
| | log.range | -0.0002 | -0.002 | 0.002 | 18,408 | 0.813 | fixed |
| | sigma | 0.008 | 0.007 | 0.008 | 16,986 | | residual |
| **Geographic range birds/mammals - fluctuations *σ*** | (Intercept) | 0.046 | 0.032 | 0.060 | 18,000 | 0.0001 | fixed |
| | log(Geographic range) | -0.001 | -0.002 | -0.001 | 18,000 | 0.001 | fixed |
| | sigma | 0.002 | 0.002 | 0.002 | 15,577 | | residual |
| **Geographic range birds/mammals - fluctuations CI** | (Intercept) | 0.180 | 0.137 | 0.226 | 18,000 | 0.0001 | fixed |
| | log(Geographic range) | -0.004 | -0.007 | -0.001 | 18,769 | 0.003 | fixed |
| | sigma | 0.011 | 0.011 | 0.011 | 18,000 | | residual |

| | | | | | | | |
|---|---|---|---|---|---|---|---|
| **Geographic range birds/mammals - fluctuations CI weighted** | (Intercept) | 0.173 | 0.117 | 0.233 | 18,000 | 0.0001 | fixed |
| | log(Geographic range) | -0.003 | -0.006 | 0.001 | 18,000 | 0.122 | fixed |
| | sigma | 0.011 | 0.010 | 0.011 | 18,000 | | residual |
| **Geographic range birds/mammals - fluctuations SE** | (Intercept) | 0.082 | 0.037 | 0.126 | 17,605 | 0.0001 | fixed |
| | log(Geographic range) | 0.001 | -0.002 | 0.004 | 17,534 | 0.512 | fixed |
| | sigma | 0.011 | 0.011 | 0.011 | 18,000 | | residual |
| **Geographic range birds/mammals - fluctuations SD** | (Intercept) | 0.605 | 0.571 | 0.638 | 17,938 | 0.0001 | fixed |
| | log(Geographic range) | -0.004 | -0.006 | -0.002 | 17,306 | 0.0004 | fixed |
| | sigma | 0.008 | 0.007 | 0.008 | 18,000 | | residual |
| **Mean population size - *μ*** | (Intercept) | 0.001 | -0.003 | 0.005 | 18,000 | 0.794 | fixed |
| | log(Mean population size) | 0.001 | 0.001 | 0.002 | 18,000 | 0.0004 | fixed |
| | sigma | 0.004 | 0.004 | 0.004 | 15,837 | | residual |
| **Mean population size - *μ* * taxa interaction** | (Intercept) | 0.007 | -0.003 | 0.018 | 18,000 | 0.161 | fixed |
| | log(Mean population size) | -0.001 | -0.003 | 0.001 | 18,000 | 0.197 | fixed |
| | Amphibia | -0.006 | -0.027 | 0.017 | 18,000 | 0.615 | fixed |
| | Aves | -0.009 | -0.021 | 0.003 | 18,000 | 0.133 | fixed |
| | Elasmobranchii | -0.025 | -0.050 | 0.0004 | 18,000 | 0.055 | fixed |
| | Mammalia | -0.009 | -0.022 | 0.006 | 18,000 | 0.211 | fixed |
| | Reptilia | 0.007 | -0.011 | 0.025 | 18,000 | 0.434 | fixed |
| | log(Mean population size):Amphibia | -0.003 | -0.008 | 0.002 | 19,145 | 0.215 | fixed |

| | | | | | | | |
|---|---|---|---|---|---|---|---|
| | log(Mean population size):Aves | 0.003 | 0.001 | 0.005 | 18,000 | 0.003 | fixed |
| | log(Mean population size):Elasmobranchii | 0.005 | -0.001 | 0.012 | 18,428 | 0.127 | fixed |
| | log(Mean population size):Mammalia | 0.004 | 0.001 | 0.006 | 18,000 | 0.002 | fixed |
| | log(Mean population size):Reptilia | 0.001 | -0.003 | 0.004 | 18,000 | 0.772 | fixed |
| | sigma | 0.004 | 0.004 | 0.004 | 16,486 | | residual |
| **Mean population size - weighted** | (Intercept) | 0.002 | -0.005 | 0.008 | 18,000 | 0.611 | fixed |
| | log(Mean population size) | 0.001 | 0.0003 | 0.002 | 18,000 | 0.014 | fixed |
| | sigma | 0.005 | 0.004 | 0.005 | 13,784 | | residual |
| **Mean population size - slope** | (Intercept) | -0.0002 | -0.007 | 0.006 | 18,000 | 0.946 | fixed |
| | log(Mean population size) | 0.002 | 0.001 | 0.003 | 18,000 | 0.0002 | fixed |
| | sigma | 0.011 | 0.010 | 0.012 | 15,825 | | residual |
| **Mean population size - fluctuations *σ*** | (Intercept) | 0.034 | 0.032 | 0.037 | 18,000 | 0.0001 | fixed |
| | log(Mean population size) | -0.001 | -0.001 | -0.0002 | 18,000 | 0.003 | fixed |
| | sigma | 0.002 | 0.002 | 0.002 | 6,763 | | residual |
| **Mean population size - fluctuations CI** | (Intercept) | 0.222 | 0.212 | 0.232 | 18,000 | 0.0001 | fixed |
| | log(Mean population size) | -0.004 | -0.006 | -0.003 | 18,000 | 0.0001 | fixed |
| | sigma | 0.026 | 0.025 | 0.027 | 14,178 | | residual |
| **Mean population size - fluctuations CI weighted** | (Intercept) | 0.262 | 0.249 | 0.275 | 18,000 | 0.0001 | fixed |

| | | | | | | | |
|---|---|---|---|---|---|---|---|
| | log(Mean population size) | -0.007 | -0.009 | -0.004 | 18,000 | 0.0001 | fixed |
| | sigma | 0.026 | 0.025 | 0.028 | 13,162 | | residual |
| **Mean population size - fluctuations SE** | (Intercept) | 0.228 | 0.218 | 0.238 | 18,000 | 0.0001 | fixed |
| | log(Mean population size) | -0.010 | -0.012 | -0.008 | 18,000 | 0.0001 | fixed |
| | sigma | 0.023 | 0.022 | 0.024 | 15,480 | | residual |
| **Mean population size - fluctuations SD** | (Intercept) | 0.001 | -0.001 | 0.003 | 17,324 | 0.454 | fixed |
| | log(Mean population size) | 0.00000 | -0.00001 | 0.00002 | 18,000 | 0.556 | fixed |
| | sigma | 0.00000 | 0.00000 | 0.00000 | 18,000 | | residual |
| **Habitat specificity - *μ*** | (Intercept) | 0.002 | -0.001 | 0.006 | 18,000 | 0.176 | fixed |
| | Habitat specificity | 0.0002 | -0.0002 | 0.001 | 18,000 | 0.321 | fixed |
| | sigma | 0.003 | 0.003 | 0.003 | 15,647 | | residual |
| **Habitat specificity - *μ* * taxa interaction** | (Intercept) | 0.003 | -0.004 | 0.011 | 18,000 | 0.443 | fixed |
| | Habitat specificity | -0.0004 | -0.002 | 0.001 | 18,187 | 0.538 | fixed |
| | Amphibia | -0.015 | -0.035 | 0.004 | 17,354 | 0.130 | fixed |
| | Aves | -0.002 | -0.011 | 0.007 | 18,000 | 0.652 | fixed |
| | Cephalaspidomorphi | 0.016 | -0.103 | 0.132 | 18,000 | 0.796 | fixed |
| | Elasmobranchii | -0.027 | -0.055 | 0.001 | 17,291 | 0.059 | fixed |
| | Holocephali | -0.105 | -0.224 | 0.013 | 18,000 | 0.082 | fixed |
| | Mammalia | 0.008 | -0.003 | 0.018 | 18,000 | 0.168 | fixed |
| | Myxini | -0.063 | -0.145 | 0.022 | 18,000 | 0.146 | fixed |
| | Reptilia | 0.015 | -0.007 | 0.036 | 18,000 | 0.176 | fixed |
| | Habitat specificity: Amphibia | 0.0003 | -0.002 | 0.002 | 17,509 | 0.753 | fixed |
| | Habitat specificity: Aves | 0.001 | -0.001 | 0.002 | 18,000 | 0.261 | fixed |
| | Habitat specificity: | 0.005 | -0.003 | 0.013 | 18,000 | 0.264 | fixed |

| | | | | | | | |
|---|---|---|---|---|---|---|---|
| | Elasmobranchii | | | | | | |
| | Habitat specificity: Mammalia | 0.001 | -0.001 | 0.002 | 19,769 | 0.494 | fixed |
| | Habitat specificity: Reptilia | -0.001 | -0.004 | 0.002 | 18,000 | 0.503 | fixed |
| | sigma | 0.003 | 0.003 | 0.003 | 16,057 | | residual |
| **Habitat specificity - weighted** | (Intercept) | 0.002 | -0.003 | 0.006 | 18,000 | 0.443 | fixed |
| | Habitat specificity | 0.0004 | -0.0001 | 0.001 | 18,000 | 0.128 | fixed |
| | sigma | 0.003 | 0.003 | 0.003 | 13,856 | | residual |
| **Habitat specificity - slope** | (Intercept) | 0.003 | -0.003 | 0.009 | 18,000 | 0.317 | fixed |
| | Habitat specificity | 0.0003 | -0.0003 | 0.001 | 18,000 | 0.321 | fixed |
| | sigma | 0.008 | 0.008 | 0.009 | 16,422 | | residual |
| **Habitat specificity - fluctuations *σ*** | (Intercept) | 0.023 | 0.021 | 0.025 | 16,596 | 0.0001 | fixed |
| | Habitat specificity | 0.0002 | -0.0001 | 0.0005 | 17,782 | 0.123 | fixed |
| | sigma | 0.002 | 0.002 | 0.002 | 14,047 | | residual |
| **Habitat specificity - fluctuations CI** | (Intercept) | 0.126 | 0.118 | 0.134 | 18,000 | 0.0001 | fixed |
| | Habitat specificity | 0.001 | -0.0002 | 0.002 | 18,000 | 0.117 | fixed |
| | sigma | 0.012 | 0.011 | 0.012 | 18,411 | | residual |
| **Habitat specificity - fluctuations CI weighted** | (Intercept) | 0.140 | 0.129 | 0.151 | 18,000 | 0.0001 | fixed |
| | Habitat specificity | 0.001 | -0.0001 | 0.002 | 18,000 | 0.065 | fixed |
| | sigma | 0.011 | 0.011 | 0.012 | 18,131 | | residual |
| **Habitat specificity - fluctuations SE** | (Intercept) | 0.117 | 0.108 | 0.126 | 18,000 | 0.0001 | fixed |
| | Habitat specificity | 0.001 | -0.0002 | 0.002 | 18,000 | 0.138 | fixed |
| | sigma | 0.012 | 0.012 | 0.012 | 17,484 | | residual |
| **Habitat specificity** | (Intercept) | 0.555 | 0.549 | 0.561 | 18,000 | 0.0001 | fixed |

| | | | | | | | |
|---|---|---|---|---|---|---|---|
| **- fluctuations SD** | | | | | | | |
| | Habitat specificity | 0.0003 | -0.0004 | 0.001 | 16,414 | 0.349 | fixed |
| | sigma | 0.008 | 0.008 | 0.008 | 18,000 | | residual |
| **IUCN Red List Categories - *μ*** | Least concern | 0.005 | 0.003 | 0.008 | 18,000 | 0.0001 | fixed |
| | Near threatened | -0.004 | -0.012 | 0.004 | 18,000 | 0.319 | fixed |
| | Vulnerable | 0.003 | -0.005 | 0.010 | 18,000 | 0.427 | fixed |
| | Endangered | -0.004 | -0.013 | 0.006 | 18,000 | 0.452 | fixed |
| | Critically endangered | -0.007 | -0.020 | 0.005 | 18,541 | 0.259 | fixed |
| | sigma | 0.003 | 0.003 | 0.003 | 15,497 | | residual |
| **IUCN Red List Categories - weighted** | Least concern | 0.007 | 0.004 | 0.010 | 18,000 | 0.0001 | fixed |
| | Near threatened | 0.001 | -0.010 | 0.011 | 18,241 | 0.852 | fixed |
| | Vulnerable | 0.003 | -0.008 | 0.013 | 18,000 | 0.626 | fixed |
| | Endangered | -0.007 | -0.020 | 0.006 | 18,000 | 0.291 | fixed |
| | Critically endangered | -0.014 | -0.032 | 0.004 | 18,000 | 0.114 | fixed |
| | sigma | 0.003 | 0.003 | 0.003 | 14,285 | | residual |
| **IUCN Red List Categories - slope** | Least concern | 0.009 | 0.005 | 0.012 | 18,905 | 0.0001 | fixed |
| | Near threatened | -0.011 | -0.024 | 0.003 | 18,000 | 0.121 | fixed |
| | Vulnerable | 0.003 | -0.010 | 0.016 | 16,528 | 0.684 | fixed |
| | Endangered | -0.007 | -0.023 | 0.009 | 18,000 | 0.423 | fixed |
| | Critically endangered | -0.013 | -0.034 | 0.009 | 18,000 | 0.254 | fixed |
| | sigma | 0.008 | 0.008 | 0.009 | 15,931 | | residual |
| **IUCN Red List Categories - fluctuations *σ*** | Least concern | 0.023 | 0.022 | 0.025 | 16,756 | 0.0001 | fixed |
| | Near threatened | 0.028 | 0.023 | 0.033 | 17,565 | 0.0001 | fixed |
| | Vulnerable | 0.027 | 0.022 | 0.032 | 20,081 | 0.0001 | fixed |

| | | | | | | | |
|---|---|---|---|---|---|---|---|
| | Endangered | 0.032 | 0.026 | 0.039 | 18,000 | 0.0001 | fixed |
| | Critically endangered | 0.039 | 0.031 | 0.048 | 18,000 | 0.0001 | fixed |
| | sigma | 0.002 | 0.002 | 0.002 | 13,906 | | residual |
| **IUCN Red List Categories - fluctuations CI** | Least concern | 0.124 | 0.118 | 0.129 | 18,000 | 0.0001 | fixed |
| | Near threatened | 0.152 | 0.134 | 0.171 | 18,000 | 0.0001 | fixed |
| | Vulnerable | 0.151 | 0.134 | 0.169 | 17,076 | 0.0001 | fixed |
| | Endangered | 0.168 | 0.146 | 0.189 | 20,476 | 0.0001 | fixed |
| | Critically endangered | 0.178 | 0.150 | 0.208 | 18,000 | 0.0001 | fixed |
| | sigma | 0.012 | 0.012 | 0.012 | 17,734 | | residual |
| **IUCN Red List Categories - fluctuations CI weighted** | Least concern | 0.140 | 0.133 | 0.147 | 18,000 | 0.0001 | fixed |
| | Near threatened | 0.172 | 0.147 | 0.196 | 17,244 | 0.0001 | fixed |
| | Vulnerable | 0.171 | 0.148 | 0.195 | 18,589 | 0.0001 | fixed |
| | Endangered | 0.194 | 0.165 | 0.222 | 18,000 | 0.0001 | fixed |
| | Critically endangered | 0.193 | 0.155 | 0.231 | 18,000 | 0.0001 | fixed |
| | sigma | 0.012 | 0.011 | 0.012 | 17,086 | | residual |
| **IUCN Red List Categories - fluctuations SE** | Least concern | 0.119 | 0.113 | 0.125 | 18,000 | 0.0001 | fixed |
| | Near threatened | 0.135 | 0.114 | 0.156 | 18,000 | 0.0001 | fixed |
| | Vulnerable | 0.127 | 0.107 | 0.147 | 18,000 | 0.0001 | fixed |
| | Endangered | 0.143 | 0.119 | 0.167 | 18,000 | 0.0001 | fixed |
| | Critically endangered | 0.149 | 0.116 | 0.180 | 17,444 | 0.0001 | fixed |
| | sigma | 0.012 | 0.012 | 0.012 | 18,000 | | residual |
| **IUCN Red List Categories -** | Least concern | 0.550 | 0.546 | 0.554 | 18,000 | 0.0001 | fixed |

| | | | | | | | |
|---|---|---|---|---|---|---|---|
| **fluctuations SD** | | | | | | | |
| | Near threatened | 0.575 | 0.561 | 0.589 | 18,000 | 0.0001 | fixed |
| | Vulnerable | 0.573 | 0.559 | 0.586 | 17,618 | 0.0001 | fixed |
| | Endangered | 0.596 | 0.580 | 0.613 | 18,000 | 0.0001 | fixed |
| | Critically endangered | 0.604 | 0.582 | 0.625 | 17,544 | 0.0001 | fixed |
| | sigma | 0.008 | 0.008 | 0.008 | 17,153 | | residual |
| **IUCN threat type - *μ*** | Fishing / harvesting aquatic resources | 0.005 | -0.0005 | 0.010 | 18,000 | 0.085 | fixed |
| | Hunting / trapping terrestrial animals | 0.005 | -0.0003 | 0.010 | 18,000 | 0.068 | fixed |
| | Annual / perennial non-timber crops | 0.005 | -0.0004 | 0.012 | 18,000 | 0.080 | fixed |
| | Intentional use (species is the target) | 0.005 | -0.001 | 0.011 | 18,000 | 0.092 | fixed |
| | Habitat shifting / alteration | 0.005 | -0.001 | 0.011 | 18,000 | 0.098 | fixed |
| | Housing / urban areas | 0.004 | -0.002 | 0.011 | 17,916 | 0.210 | fixed |
| | Unintentional effects: (large scale) [harvest] | 0.004 | -0.002 | 0.010 | 19,228 | 0.205 | fixed |
| | Intentional use: (subsistence/small scale) [harvest] | 0.005 | -0.002 | 0.012 | 18,423 | 0.192 | fixed |
| | Agricultural / forestry effluents | 0.005 | -0.002 | 0.011 | 18,000 | 0.153 | fixed |
| | Intentional use: (large scale) [harvest] | 0.004 | -0.003 | 0.011 | 17,575 | 0.257 | fixed |
| | sigma | 0.004 | 0.004 | 0.004 | 18,000 | | residual |
| **IUCN threat number - *μ*** | (Intercept) | 0.004 | -0.001 | 0.009 | 18,000 | 0.145 | fixed |
| | Number of threats | 0.0001 | -0.0004 | 0.001 | 18,000 | 0.726 | fixed |

| | | | | | | |
|---|---|---|---|---|---|---|
| | sigma | 0.004 | 0.004 | 0.004 | 16,084 | | residual |
| **IUCN threat type - fluctuations *σ*** | Fishing / harvesting aquatic resources | 0.033 | 0.028 | 0.037 | 17,468 | 0.0001 | fixed |
| | Hunting / trapping terrestrial animals | 0.033 | 0.029 | 0.038 | 17,547 | 0.0001 | fixed |
| | Annual / perennial non-timber crops | 0.033 | 0.029 | 0.038 | 17,225 | 0.0001 | fixed |
| | Intentional use (species is the target) | 0.033 | 0.029 | 0.038 | 18,000 | 0.0001 | fixed |
| | Habitat shifting / alteration | 0.032 | 0.028 | 0.037 | 17,548 | 0.0001 | fixed |
| | Housing / urban areas | 0.034 | 0.029 | 0.039 | 18,000 | 0.0001 | fixed |
| | Unintentional effects: (large scale) [harvest] | 0.033 | 0.028 | 0.037 | 17,134 | 0.0001 | fixed |
| | Intentional use: (subsistence/small scale) [harvest] | 0.033 | 0.028 | 0.038 | 18,000 | 0.0001 | fixed |
| | Agricultural / forestry effluents | 0.033 | 0.028 | 0.037 | 17,571 | 0.0001 | fixed |
| | Intentional use: (large scale) [harvest] | 0.032 | 0.027 | 0.037 | 18,000 | 0.0001 | fixed |
| | sigma | 0.002 | 0.002 | 0.002 | 18,000 | | residual |
| **IUCN threat number - fluctuations *σ*** | (Intercept) | 0.029 | 0.026 | 0.032 | 17,078 | 0.0001 | fixed |
| | Number of threats | 0.0001 | -0.0002 | 0.0004 | 18,000 | 0.475 | fixed |
| | sigma | 0.002 | 0.002 | 0.002 | 7,021 | | residual |

**Table S3. Model outputs from UK-scale analyses.** Sigma is the overall model residual variance. Net population change is estimated using $\mu$ values derived from state-space models of population abundance versus time and slopes of linear models of population abundance versus time. The weighted $\mu$ models included $\mu$ as a response variable, weighted by $\tau$, the observation error estimate derived from the state-space models. The fluctuation models included the process noise ($\sigma^2$) values from state-space models, half of the 95% confidence interval around the $\mu$ value of population change, the standard error around the slopes of linear models of population abundance versus time, and the standard deviation of the raw time-series data for each population. The process noise is a metric of population fluctuations, whereas the rest of the metrics show population variability. The weighted fluctuation models were weighted by $\tau$, the observation error estimate derived from the state-space models.

| Model name | Variable | Posterior mean | Lower 95% CI | Upper 95% CI | Effective sample size | pMCMC | Effect |
|---|---|---|---|---|---|---|---|
| **Realm - $\mu$** | Terrestrial | 0.011 | -0.003 | 0.023 | 18,000 | 0.105 | fixed |
| | Marine | 0.007 | -0.006 | 0.020 | 18,000 | 0.297 | fixed |
| | Freshwater | 0.030 | 0.010 | 0.049 | 17,098 | 0.002 | fixed |
| | sigma | 0.003 | 0.003 | 0.004 | 13,565 | | residual |
| **Realm - weighted** | Terrestrial | 0.015 | 0.0004 | 0.030 | 18,000 | 0.038 | fixed |
| | Marine | 0.008 | -0.008 | 0.024 | 18,000 | 0.303 | fixed |
| | Freshwater | 0.027 | 0.005 | 0.050 | 18,000 | 0.021 | fixed |
| | sigma | 0.003 | 0.002 | 0.004 | 10,568 | | residual |
| **Realm - slope** | Freshwater | 0.062 | 0.025 | 0.098 | 18,000 | 0.0003 | fixed |
| | Marine | 0.007 | -0.018 | 0.031 | 18,000 | 0.601 | fixed |
| | Terrestrial | 0.020 | -0.004 | 0.045 | 18,380 | 0.114 | fixed |
| | sigma | 0.011 | 0.009 | 0.012 | 15,525 | | residual |
| **Realm - fluctuations $\sigma$** | Terrestrial | 0.085 | 0.062 | 0.105 | 18,000 | 0.0001 | fixed |
| | Marine | 0.056 | 0.034 | 0.079 | 18,000 | 0.0001 | fixed |
| | Freshwater | 0.051 | 0.019 | 0.083 | 18,000 | 0.002 | fixed |
| | sigma | 0.005 | 0.004 | 0.006 | 16,954 | | residual |
| **Realm - fluctuations CI** | Terrestrial | 0.193 | 0.165 | 0.220 | 18,000 | 0.0001 | fixed |
| | Marine | 0.118 | 0.090 | 0.146 | 18,000 | 0.0001 | fixed |
| | Freshwater | 0.139 | 0.100 | 0.181 | 18,000 | 0.0001 | fixed |
| | sigma | 0.011 | 0.009 | 0.013 | 15,777 | | residual |
| **Realm - fluctuations CI weighted** | Terrestrial | 0.191 | 0.157 | 0.224 | 18,000 | 0.0001 | fixed |

| | | | | | | | |
|---|---|---|---|---|---|---|---|
| | Marine | 0.134 | 0.098 | 0.172 | 18,000 | 0.0001 | fixed |
| | Freshwater | 0.164 | 0.115 | 0.217 | 18,001 | 0.0001 | fixed |
| | sigma | 0.010 | 0.007 | 0.012 | 15,014 | | residual |
| **Taxa - $\mu$** | Actinopterygii | -0.003 | -0.020 | 0.014 | 16,879 | 0.744 | fixed |
| | Amphibia | -0.0002 | -0.051 | 0.052 | 18,000 | 0.995 | fixed |
| | Aves | 0.012 | 0.001 | 0.022 | 18,000 | 0.027 | fixed |
| | Elasmobranchii | 0.006 | -0.045 | 0.056 | 17,998 | 0.811 | fixed |
| | Mammalia | 0.046 | 0.023 | 0.069 | 17,742 | 0.0001 | fixed |
| | Reptilia | 0.044 | -0.018 | 0.108 | 18,000 | 0.181 | fixed |
| | sigma | 0.004 | 0.003 | 0.004 | 10,343 | | residual |
| **Taxa - weighted** | Actinopterygii | -0.010 | -0.033 | 0.011 | 18,000 | 0.374 | fixed |
| | Amphibia | -0.002 | -0.065 | 0.061 | 18,000 | 0.947 | fixed |
| | Aves | 0.017 | 0.005 | 0.029 | 18,000 | 0.005 | fixed |
| | Elasmobranchii | 0.002 | -0.089 | 0.086 | 18,000 | 0.972 | fixed |
| | Mammalia | 0.043 | 0.015 | 0.069 | 18,000 | 0.002 | fixed |
| | Reptilia | 0.030 | -0.039 | 0.103 | 18,000 | 0.404 | fixed |
| | sigma | 0.003 | 0.002 | 0.004 | 9,676 | | residual |
| **Taxa - slope** | Actinopterygii | -0.009 | -0.042 | 0.024 | 17,082 | 0.593 | fixed |
| | Amphibia | 0.004 | -0.095 | 0.111 | 18,000 | 0.931 | fixed |
| | Aves | 0.020 | 0.0001 | 0.040 | 18,434 | 0.046 | fixed |
| | Elasmobranchii | -0.014 | -0.106 | 0.078 | 18,000 | 0.766 | fixed |
| | Mammalia | 0.099 | 0.053 | 0.145 | 18,000 | 0.0001 | fixed |
| | Reptilia | 0.069 | -0.048 | 0.185 | 19,449 | 0.241 | fixed |
| | sigma | 0.011 | 0.009 | 0.013 | 14,664 | | residual |
| **Taxa - fluctuations $\sigma$** | Actinopterygii | 0.051 | 0.019 | 0.082 | 18,000 | 0.001 | fixed |
| | Amphibia | 0.083 | -0.023 | 0.182 | 18,000 | 0.117 | fixed |
| | Aves | 0.073 | 0.053 | 0.091 | 18,000 | 0.0001 | fixed |
| | Elasmobranchii | 0.072 | -0.010 | 0.154 | 18,000 | 0.083 | fixed |
| | Mammalia | 0.054 | 0.012 | 0.098 | 18,000 | 0.014 | fixed |
| | Reptilia | 0.123 | 0.014 | 0.222 | 18,000 | 0.021 | fixed |
| | sigma | 0.005 | 0.004 | 0.006 | 17,207 | | residual |
| **Taxa - fluctuations CI** | Actinopterygii | 0.113 | 0.074 | 0.153 | 18,891 | 0.0001 | fixed |
| | Amphibia | 0.230 | 0.107 | 0.358 | 18,000 | 0.0004 | fixed |
| | Aves | 0.166 | 0.143 | 0.190 | 18,000 | 0.0001 | fixed |
| | Elasmobranchii | 0.186 | 0.078 | 0.290 | 18,000 | 0.001 | fixed |
| | Mammalia | 0.150 | 0.095 | 0.204 | 18,761 | 0.0001 | fixed |
| | Reptilia | 0.275 | 0.146 | 0.410 | 18,000 | 0.0002 | fixed |
| | sigma | 0.011 | 0.009 | 0.013 | 15,974 | | residual |
| **Taxa - fluctuations CI weighted** | Actinopterygii | 0.138 | 0.088 | 0.190 | 18,000 | 0.0001 | fixed |
| | Amphibia | 0.287 | 0.134 | 0.448 | 18,000 | 0.001 | fixed |
| | Aves | 0.169 | 0.142 | 0.197 | 18,255 | 0.0001 | fixed |
| | Elasmobranchii | 0.266 | 0.097 | 0.432 | 17,623 | 0.003 | fixed |
| | Mammalia | 0.156 | 0.090 | 0.222 | 18,000 | 0.0001 | fixed |
| | Reptilia | 0.311 | 0.158 | 0.474 | 18,000 | 0.0001 | fixed |

| | | | | | | | |
|---|---|---|---|---|---|---|---|
| | sigma | 0.009 | 0.007 | 0.012 | 14,900 | | residual |
| **Geographic range (all) - *μ*** | (Intercept) | 0.006 | -0.052 | 0.063 | 18,000 | 0.850 | fixed |
| | log(km2_range) | 0.001 | -0.003 | 0.005 | 18,000 | 0.762 | fixed |
| | sigma | 0.004 | 0.003 | 0.004 | 14,513 | | residual |
| **Geographic range (all) - weighted** | (Intercept) | -0.012 | -0.077 | 0.050 | 18,000 | 0.710 | fixed |
| | log.range | 0.002 | -0.002 | 0.006 | 18,000 | 0.346 | fixed |
| | sigma | 0.003 | 0.002 | 0.004 | 13,236 | | residual |
| **Geographic range (all) - slope** | (Intercept) | 0.009 | -0.105 | 0.120 | 18,239 | 0.870 | fixed |
| | log.range | 0.001 | -0.006 | 0.009 | 18,257 | 0.755 | fixed |
| | sigma | 0.011 | 0.009 | 0.013 | 16,452 | | residual |
| **Geographic range (all) - fluctuations *σ*** | (Intercept) | 0.004 | -0.101 | 0.115 | 18,000 | 0.939 | fixed |
| | log(km2_range) | 0.005 | -0.003 | 0.012 | 18,000 | 0.198 | fixed |
| | sigma | 0.005 | 0.004 | 0.006 | 18,000 | | residual |
| **Geographic range (all) - fluctuations CI** | (Intercept) | 0.045 | -0.086 | 0.177 | 18,000 | 0.494 | fixed |
| | log.range | 0.008 | -0.001 | 0.017 | 18,000 | 0.079 | fixed |
| | sigma | 0.011 | 0.009 | 0.013 | 16,713 | | residual |
| **Geographic range (all) - fluctuations CI weighted** | (Intercept) | 0.059 | -0.104 | 0.221 | 18,000 | 0.472 | fixed |
| | log.range | 0.008 | -0.003 | 0.019 | 18,000 | 0.162 | fixed |
| | sigma | 0.010 | 0.008 | 0.013 | 15,715 | | residual |
| **Geographic range (all) - fluctuations SE** | (Intercept) | 0.050 | -0.104 | 0.206 | 18,000 | 0.528 | fixed |
| | log.range | 0.008 | -0.003 | 0.018 | 18,000 | 0.159 | fixed |
| | sigma | 0.009 | 0.007 | 0.010 | 18,000 | | residual |
| **Geographic range (all) - fluctuations SD** | (Intercept) | 0.171 | 0.039 | 0.294 | 18,000 | 0.008 | fixed |
| | log.range | 0.016 | 0.007 | 0.025 | 18,000 | 0.0002 | fixed |
| | sigma | 0.017 | 0.014 | 0.020 | 15,395 | | residual |
| **Mean population size - *μ*** | (Intercept) | 0.023 | 0.0001 | 0.047 | 1,207 | 0.051 | fixed |
| | log(Mean population size) | -0.0004 | -0.004 | 0.004 | 2,374 | 0.860 | fixed |
| | sigma | 0.006 | 0.005 | 0.008 | 411 | | residual |
| **Mean population size - weighted** | (Intercept) | 0.021 | -0.002 | 0.045 | 852 | 0.066 | fixed |
| | log(Mean population size) | -0.0001 | -0.004 | 0.004 | 1,950 | 0.964 | fixed |
| | sigma | 0.007 | 0.005 | 0.009 | 222 | | residual |
| **Mean population size - slope** | (Intercept) | 0.035 | -0.006 | 0.075 | 1,274 | 0.071 | fixed |
| | log(meanpop) | -0.001 | -0.008 | 0.006 | 6,788 | 0.832 | fixed |

| | | | | | | | |
|---|---|---|---|---|---|---|---|
| | sigma | 0.022 | 0.016 | 0.027 | 593 | | residual |
| **Mean population size - fluctuations *σ*** | (Intercept) | 0.169 | 0.118 | 0.216 | 18,000 | 0.0001 | fixed |
| | log(Mean population size) | -0.004 | -0.013 | 0.005 | 18,000 | 0.389 | fixed |
| | sigma | 0.034 | 0.023 | 0.045 | 200 | | residual |
| **Mean population size - fluctuations CI** | (Intercept) | 0.298 | 0.242 | 0.357 | 18,000 | 0.0001 | fixed |
| | log(Mean population size) | -0.009 | -0.019 | 0.001 | 18,000 | 0.080 | fixed |
| | sigma | 0.026 | 0.018 | 0.035 | 11,283 | | residual |
| **Mean population size - fluctuations CI weighted** | (Intercept) | 0.298 | 0.240 | 0.354 | 18,305 | 0.0001 | fixed |
| | log(Mean population size) | -0.009 | -0.019 | 0.001 | 18,000 | 0.077 | fixed |
| | sigma | 0.026 | 0.018 | 0.035 | 10,953 | | residual |
| **Mean population size - fluctuations SE** | (Intercept) | 0.354 | 0.293 | 0.411 | 18,000 | 0.0001 | fixed |
| | log(meanpop) | -0.024 | -0.034 | -0.014 | 18,099 | 0.0001 | fixed |
| | sigma | 0.014 | 0.009 | 0.018 | 16,950 | | residual |
| **Mean population size - fluctuations SD** | (Intercept) | 0.519 | 0.473 | 0.565 | 17,529 | 0.0001 | fixed |
| | log(Mean population size) | -0.007 | -0.016 | 0.001 | 17,123 | 0.086 | fixed |
| | sigma | 0.030 | 0.019 | 0.039 | 83 | | residual |
| **Habitat specificity - *μ*** | (Intercept) | 0.007 | -0.010 | 0.024 | 18,000 | 0.449 | fixed |
| | Habitat specificity | 0.001 | -0.001 | 0.002 | 18,000 | 0.340 | fixed |
| | sigma | 0.004 | 0.003 | 0.005 | 13,436 | | residual |
| **Habitat specificity (profiling) - *μ*** | (Intercept) | 0.013 | -0.005 | 0.030 | 18,000 | 0.173 | fixed |
| | Habitat specificity | -0.0002 | -0.003 | 0.002 | 18,000 | 0.864 | fixed |
| | sigma | 0.005 | 0.004 | 0.005 | 8,201 | | residual |
| **Habitat specificity - weighted** | (Intercept) | 0.009 | -0.012 | 0.030 | 18,441 | 0.381 | fixed |
| | Habitat specificity | 0.001 | -0.001 | 0.003 | 18,000 | 0.424 | fixed |
| | sigma | 0.003 | 0.002 | 0.004 | 10,360 | | residual |
| **Habitat specificity - slope** | (Intercept) | 0.013 | -0.020 | 0.046 | 17,956 | 0.434 | fixed |
| | Habitat specificity | 0.001 | -0.002 | 0.004 | 17,589 | 0.434 | fixed |
| | sigma | 0.011 | 0.009 | 0.014 | 13,969 | | residual |
| **Habitat specificity (profiling) - slope** | (Intercept) | 0.015 | -0.020 | 0.049 | 18,509 | 0.394 | fixed |

| | | | | | | | |
|---|---|---|---|---|---|---|---|
| | Habitat specificity | 0.0004 | -0.004 | 0.005 | 17,454 | 0.852 | fixed |
| | sigma | 0.013 | 0.011 | 0.016 | 12,446 | | residual |
| **Habitat specificity - fluctuations *σ*** | (Intercept) | 0.055 | 0.027 | 0.085 | 18,000 | 0.0003 | fixed |
| | Habitat specificity | 0.002 | -0.001 | 0.004 | 17,598 | 0.221 | fixed |
| | sigma | 0.005 | 0.004 | 0.006 | 17,516 | | residual |
| **Habitat specificity (profiling) - fluctuations *σ*** | (Intercept) | 0.022 | 0.008 | 0.038 | 18,947 | 0.004 | fixed |
| | Habitat specificity | 0.002 | -0.0003 | 0.004 | 18,000 | 0.092 | fixed |
| | sigma | 0.002 | 0.001 | 0.002 | 12,444 | | residual |
| **Habitat specificity - fluctuations CI** | (Intercept) | 0.138 | 0.101 | 0.176 | 18,800 | 0.0001 | fixed |
| | Habitat specificity | 0.003 | -0.001 | 0.006 | 18,000 | 0.137 | fixed |
| | sigma | 0.011 | 0.009 | 0.014 | 15,996 | | residual |
| **Habitat specificity - fluctuations CI weighted** | (Intercept) | 0.146 | 0.098 | 0.192 | 18,046 | 0.0001 | fixed |
| | Habitat specificity | 0.003 | -0.001 | 0.007 | 19,218 | 0.205 | fixed |
| | sigma | 0.010 | 0.008 | 0.013 | 15,033 | | residual |
| **Habitat specificity - fluctuations SE** | (Intercept) | 0.155 | 0.114 | 0.198 | 18,000 | 0.0001 | fixed |
| | Habitat specificity | 0.001 | -0.003 | 0.004 | 18,000 | 0.772 | fixed |
| | sigma | 0.009 | 0.007 | 0.010 | 18,000 | | residual |
| **Habitat specificity - fluctuations SD** | (Intercept) | 0.400 | 0.360 | 0.438 | 17,592 | 0.0001 | fixed |
| | Habitat specificity | 0.0004 | -0.003 | 0.004 | 18,000 | 0.844 | fixed |
| | sigma | 0.017 | 0.014 | 0.020 | 15,211 | | residual |
| **IUCN Red List Categories - *μ*** | Least concern | 0.013 | 0.003 | 0.023 | 17,332 | 0.011 | fixed |
| | Near threatened | 0.020 | -0.014 | 0.053 | 18,436 | 0.259 | fixed |
| | Vulnerable | 0.014 | -0.028 | 0.059 | 18,000 | 0.516 | fixed |
| | Endangered | -0.038 | -0.186 | 0.114 | 18,000 | 0.621 | fixed |
| | Critically endangered | 0.070 | -0.024 | 0.166 | 17,353 | 0.148 | fixed |
| | sigma | 0.004 | 0.003 | 0.004 | 14,117 | | residual |
| **IUCN Red List Categories - weighted** | Least concern | 0.017 | 0.006 | 0.029 | 17,546 | 0.003 | fixed |
| | Near threatened | 0.025 | -0.017 | 0.064 | 16,702 | 0.222 | fixed |
| | Vulnerable | 0.001 | -0.046 | 0.048 | 18,000 | 0.964 | fixed |
| | Endangered | -0.038 | -0.181 | 0.102 | 18,000 | 0.598 | fixed |
| | Critically endangered | 0.044 | -0.200 | 0.283 | 16,964 | 0.713 | fixed |
| | sigma | 0.003 | 0.002 | 0.004 | 10,497 | | residual |

| | | | | | | | |
|---|---|---|---|---|---|---|---|
| **IUCN Red List Categories - slope** | Least concern | 0.026 | 0.007 | 0.046 | 18,000 | 0.009 | fixed |
| | Near threatened | 0.016 | -0.053 | 0.079 | 18,000 | 0.635 | fixed |
| | Vulnerable | 0.017 | -0.069 | 0.102 | 18,000 | 0.689 | fixed |
| | Endangered | -0.011 | -0.289 | 0.275 | 18,091 | 0.934 | fixed |
| | Critically endangered | 0.053 | -0.129 | 0.235 | 18,475 | 0.566 | fixed |
| | sigma | 0.011 | 0.009 | 0.013 | 15,903 | | residual |
| **IUCN Red List Categories - fluctuations** $\sigma$ | Least concern | 0.073 | 0.056 | 0.090 | 19,089 | 0.0001 | fixed |
| | Near threatened | 0.052 | -0.012 | 0.113 | 18,000 | 0.099 | fixed |
| | Vulnerable | 0.061 | -0.018 | 0.141 | 18,461 | 0.139 | fixed |
| | Endangered | 0.173 | -0.072 | 0.417 | 18,000 | 0.165 | fixed |
| | Critically endangered | 0.003 | -0.159 | 0.166 | 18,000 | 0.976 | fixed |
| | sigma | 0.005 | 0.004 | 0.006 | 17,270 | | residual |
| **IUCN Red List Categories - fluctuations CI** | Least concern | 0.165 | 0.144 | 0.187 | 18,000 | 0.0001 | fixed |
| | Near threatened | 0.134 | 0.057 | 0.211 | 18,000 | 0.001 | fixed |
| | Vulnerable | 0.140 | 0.041 | 0.240 | 18,000 | 0.005 | fixed |
| | Endangered | 0.237 | -0.089 | 0.541 | 18,433 | 0.140 | fixed |
| | Critically endangered | 0.105 | -0.103 | 0.314 | 17,573 | 0.321 | fixed |
| | sigma | 0.011 | 0.009 | 0.013 | 16,778 | | residual |
| **IUCN Red List Categories - fluctuations CI weighted** | Least concern | 0.175 | 0.148 | 0.201 | 18,000 | 0.0001 | fixed |
| | Near threatened | 0.166 | 0.071 | 0.263 | 18,000 | 0.001 | fixed |
| | Vulnerable | 0.148 | 0.034 | 0.264 | 18,000 | 0.013 | fixed |
| | Endangered | 0.236 | -0.100 | 0.568 | 18,000 | 0.176 | fixed |
| | Critically endangered | 0.070 | -0.256 | 0.420 | 18,493 | 0.690 | fixed |
| | sigma | 0.010 | 0.008 | 0.012 | 15,655 | | residual |
| **IUCN Red List Categories - fluctuations SE** | Least concern | 0.165 | 0.141 | 0.190 | 18,376 | 0.0001 | fixed |
| | Near threatened | 0.134 | 0.048 | 0.227 | 14,776 | 0.004 | fixed |
| | Vulnerable | 0.113 | -0.002 | 0.228 | 17,971 | 0.051 | fixed |
| | Endangered | 0.071 | -0.281 | 0.409 | 18,000 | 0.692 | fixed |
| | Critically endangered | 0.079 | -0.156 | 0.315 | 18,000 | 0.516 | fixed |
| | sigma | 0.009 | 0.007 | 0.010 | 18,000 | | residual |
| **IUCN Red List Categories - fluctuations SD** | Least concern | 0.408 | 0.386 | 0.429 | 18,770 | 0.0001 | fixed |
| | Near threatened | 0.358 | 0.287 | 0.432 | 18,000 | 0.0001 | fixed |
| | Vulnerable | 0.319 | 0.223 | 0.418 | 18,000 | 0.0001 | fixed |
| | Endangered | 0.243 | -0.085 | 0.586 | 18,000 | 0.157 | fixed |

| | | | | | | | |
|---|---|---|---|---|---|---|---|
| | Critically endangered | 0.331 | 0.129 | 0.536 | 18,000 | 0.002 | fixed |
| | sigma | 0.016 | 0.015 | 0.016 | 18,000 | | residual |

**Table S4. Phylogeny model outputs.** To account for phylogenetic uncertainty, we ran the phylogenetic models for amphibian, bird and reptile species using 10 different trees for each class, and here we present the mean, min and max values from the different model runs. Sigma is the overall model residual variance. Net population change is estimated using $\mu$ values derived from state-space models of population abundance versus time and slopes of linear models of population abundance versus time. The fluctuation models were based on the process noise ($\sigma^2$) values from state-space models.

| **Model name** | **Variable** | **Mean pMCMC** | **Max pMCMC** | **Min pMCMC** | **Mean effective sample size** | **Mean post. mean** | **Max post. mean** | **Min post. mean** | **Mean lower 95% CI** | **Max lower 95% CI** | **Min lower 95% CI** | **Mean upper 95% CI** | **Max upper 95% CI** | **Min upper 95% CI** |
|---|---|---|---|---|---|---|---|---|---|---|---|---|---|---|
| **Amphibian population trends** | (Intercept) | 0.587 | 0.621 | 0.558 | 10047 | -0.009 | -0.008 | -0.009 | -0.051 | -0.046 | -0.054 | 0.031 | 0.035 | 0.027 |
| | Phylogeny | | | | 7037 | 0.001 | 0.001 | 0.001 | 0 | 0 | 0 | 0.004 | 0.005 | 0.004 |
| | Sigma | | | | 9856 | 0.006 | 0.006 | 0.006 | 0.004 | 0.004 | 0.004 | 0.007 | 0.007 | 0.007 |
| | Species | | | | 7912 | 0.001 | 0.001 | 0 | 0 | 0 | 0 | 0.002 | 0.002 | 0.002 |
| **Amphibian population fluctuations** | (Intercept) | 0 | 0 | 0 | 9947 | 0.155 | 0.156 | 0.155 | 0.104 | 0.105 | 0.102 | 0.208 | 0.211 | 0.206 |
| | Phylogeny | | | | 9778 | 0.001 | 0.001 | 0.001 | 0 | 0 | 0 | 0.005 | 0.006 | 0.005 |
| | Sigma | | | | 10172 | 0.048 | 0.048 | 0.048 | 0.037 | 0.038 | 0.037 | 0.059 | 0.06 | 0.059 |
| | Species | | | | 9817 | 0.001 | 0.001 | 0.001 | 0 | 0 | 0 | 0.003 | 0.003 | 0.003 |
| **Bird population trends** | (Intercept) | 0.449 | 0.63 | 0.305 | 10233 | 0.005 | 0.005 | 0.004 | -0.009 | -0.005 | -0.014 | 0.018 | 0.021 | 0.016 |
| | Phylogeny | | | | 3770 | 0 | 0 | 0 | 0 | 0 | 0 | 0 | 0.001 | 0 |
| | Sigma | | | | 9992 | 0.002 | 0.002 | 0.002 | 0.002 | 0.002 | 0.002 | 0.003 | 0.003 | 0.003 |
| | Species | | | | 6682 | 0 | 0 | 0 | 0 | 0 | 0 | 0 | 0 | 0 |
| **Bird population fluctuations** | (Intercept) | 0.001 | 0.006 | 0 | 10183 | 0.02 | 0.02 | 0.019 | 0.01 | 0.013 | 0.007 | 0.029 | 0.032 | 0.027 |
| | Phylogeny | | | | 3995 | 0 | 0 | 0 | 0 | 0 | 0 | 0 | 0 | 0 |

| | | | | | | | | | | | | | | |
|---|---|---|---|---|---|---|---|---|---|---|---|---|---|---|
| | Sigma | | | | 9221 | 0.002 | 0.002 | 0.002 | 0.002 | 0.002 | 0.002 | 0.002 | 0.002 | 0.002 |
| | Species | | | | 5322 | 0 | 0 | 0 | 0 | 0 | 0 | 0 | 0 | 0 |
| **Reptile population trends** | (Intercept) | 0.832 | 0.856 | 0.812 | 10030 | 0.004 | 0.005 | 0.003 | -0.038 | -0.033 | -0.047 | 0.048 | 0.056 | 0.043 |
| | Phylogeny | | | | 3971 | 0.002 | 0.002 | 0.001 | 0 | 0 | 0 | 0.007 | 0.01 | 0.005 |
| | Sigma | | | | 3562 | 0.004 | 0.004 | 0.004 | 0.001 | 0.001 | 0.001 | 0.007 | 0.007 | 0.007 |
| | Species | | | | 3419 | 0.004 | 0.004 | 0.004 | 0 | 0 | 0 | 0.007 | 0.007 | 0.007 |
| **Reptile population fluctuations** | (Intercept) | 0.003 | 0.004 | 0.001 | 9839 | 0.151 | 0.152 | 0.151 | 0.074 | 0.078 | 0.07 | 0.23 | 0.236 | 0.225 |
| | Phylogeny | | | | 2958 | 0.004 | 0.005 | 0.004 | 0 | 0 | 0 | 0.017 | 0.019 | 0.016 |
| | Sigma | | | | 3888 | 0.007 | 0.007 | 0.007 | 0.003 | 0.003 | 0.002 | 0.014 | 0.015 | 0.014 |
| | Species | | | | 6057 | 0.043 | 0.043 | 0.043 | 0.025 | 0.026 | 0.024 | 0.062 | 0.064 | 0.061 |

**Table S5. List of species included in the UK scale analysis of population change across rarity metrics.**

| Species name | Number of populations |
|---|---|
| *Acrocephalus schoenobaenus* | 1 |
| *Acrocephalus scirpaceus* | 1 |
| *Agonus cataphractus* | 1 |
| *Alca torda* | 4 |
| *Anarhichas lupus* | 1 |
| *Anas acuta* | 1 |
| *Anas crecca* | 1 |
| *Anas platyrhynchos* | 2 |
| *Anser albifrons* | 4 |
| *Anser fabalis* | 1 |
| *Anthus pratensis* | 2 |
| *Ardea cinerea* | 1 |
| *Arenaria interpres* | 1 |
| *Argentina silus* | 1 |
| *Argentina sphyraena* | 1 |
| *Arnoglossus laterna* | 1 |
| *Asio flammeus* | 1 |
| *Aythya ferina* | 4 |
| *Aythya fuligula* | 2 |
| *Botaurus stellaris* | 14 |
| *Branta bernicla* | 54 |
| *Branta canadensis* | 1 |
| *Branta leucopsis* | 3 |
| *Brosme brosme* | 1 |
| *Bucephala clangula* | 1 |
| *Bufo bufo* | 1 |
| *Burhinus oedicnemus* | 1 |
| *Buteo buteo* | 1 |
| *Calidris alba* | 1 |
| *Calidris alpina* | 1 |
| *Calidris canutus* | 2 |
| *Calidris maritima* | 2 |
| *Callionymus maculatus* | 1 |
| *Capreolus capreolus* | 3 |

| | |
|---|---|
| *Carduelis cannabina* | 2 |
| *Cepphus grylle* | 2 |
| *Cervus elaphus* | 2 |
| *Cetorhinus maximus* | 1 |
| *Cettia cetti* | 1 |
| *Charadrius hiaticula* | 1 |
| *Chelidonichthys lucerna* | 2 |
| *Circus aeruginosus* | 1 |
| *Circus cyaneus* | 1 |
| *Clupea harengus* | 2 |
| *Columba oenas* | 1 |
| *Coronella austriaca* | 1 |
| *Corvus corax* | 1 |
| *Corvus corone* | 1 |
| *Corvus monedula* | 1 |
| *Crex crex* | 1 |
| *Cyclopterus lumpus* | 1 |
| *Cygnus columbianus* | 3 |
| *Cygnus cygnus* | 1 |
| *Cygnus olor* | 1 |
| *Delphinus delphis* | 2 |
| *Echiichthys vipera* | 1 |
| *Egretta garzetta* | 1 |
| *Emberiza cirlus* | 1 |
| *Emberiza citrinella* | 1 |
| *Emberiza schoeniclus* | 3 |
| *Eptesicus serotinus* | 1 |
| *Esox lucius* | 2 |
| *Falco peregrinus* | 1 |
| *Falco tinnunculus* | 1 |
| *Fulica atra* | 1 |
| *Fulmarus glacialis* | 7 |
| *Gadus morhua* | 7 |
| *Glyptocephalus cynoglossus* | 1 |
| *Haematopus ostralegus* | 2 |
| *Haliaeetus albicilla* | 1 |
| *Halichoerus grypus* | 52 |
| *Hippoglossus hippoglossus* | 1 |

| | |
|---|---|
| *Lagopus lagopus* | 3 |
| *Larus argentatus* | 1 |
| *Larus canus* | 1 |
| *Larus fuscus* | 1 |
| *Larus melanocephalus* | 1 |
| *Lepidorhombus whiffiagonis* | 2 |
| *Lepus timidus* | 1 |
| *Limosa lapponica* | 2 |
| *Limosa limosa* | 2 |
| *Lissotriton vulgaris* | 5 |
| *Lophius budegassa* | 1 |
| *Lophius piscatorius* | 2 |
| *Lullula arborea* | 1 |
| *Melanogrammus aeglefinus* | 9 |
| *Meles meles* | 1 |
| *Mergus serrator* | 1 |
| *Merlangius merlangus* | 8 |
| *Merluccius merluccius* | 2 |
| *Micromesistius poutassou* | 1 |
| *Milvus milvus* | 3 |
| *Molva molva* | 1 |
| *Morus bassanus* | 4 |
| *Muscicapa striata* | 1 |
| *Myotis nattereri* | 1 |
| *Natrix natrix* | 1 |
| *Netta rufina* | 1 |
| *Numenius arquata* | 2 |
| *Nyctalus noctula* | 1 |
| *Oenanthe oenanthe* | 1 |
| *Orcinus orca* | 1 |
| *Oriolus oriolus* | 1 |
| *Oryctolagus cuniculus* | 6 |
| *Oxyura jamaicensis* | 2 |
| *Pandion haliaetus* | 1 |
| *Parus major* | 2 |
| *Passer domesticus* | 1 |
| *Passer montanus* | 1 |
| *Perca fluviatilis* | 3 |

| | |
|---|---|
| *Perdix perdix* | 1 |
| *Phalacrocorax aristotelis* | 8 |
| *Phoca vitulina* | 28 |
| *Phrynorhombus norvegicus* | 1 |
| *Pipistrellus pipistrellus* | 1 |
| *Pipistrellus pygmaeus* | 1 |
| *Platichthys flesus* | 1 |
| *Plecotus auritus* | 1 |
| *Plectrophenax nivalis* | 2 |
| *Pleuronectes platessa* | 6 |
| *Pluvialis apricaria* | 2 |
| *Pluvialis squatarola* | 1 |
| *Podiceps cristatus* | 1 |
| *Pollachius pollachius* | 1 |
| *Pollachius virens* | 3 |
| *Prunella modularis* | 1 |
| *Puffinus mauretanicus* | 2 |
| *Pyrrhula pyrrhula* | 1 |
| *Raja brachyura* | 1 |
| *Raja clavata* | 1 |
| *Raja microocellata* | 1 |
| *Raja montagui* | 1 |
| *Rhinolophus ferrumequinum* | 2 |
| *Rhinolophus hipposideros* | 4 |
| *Rissa tridactyla* | 9 |
| *Salmo salar* | 1 |
| *Salmo trutta* | 1 |
| *Scomber scombrus* | 2 |
| *Scyliorhinus canicula* | 1 |
| *Sitta europaea* | 1 |
| *Sprattus sprattus* | 1 |
| *Stenella coeruleoalba* | 1 |
| *Stercorarius parasiticus* | 3 |
| *Sterna dougallii* | 5 |
| *Sterna hirundo* | 1 |
| *Sterna paradisaea* | 1 |
| *Sternula albifrons* | 1 |
| *Streptopelia turtur* | 1 |

| | |
|---|---|
| *Strix aluco* | 1 |
| *Sturnus vulgaris* | 1 |
| *Sylvia communis* | 1 |
| *Syngnathus rostellatus* | 1 |
| *Tachybaptus ruficollis* | 1 |
| *Tadorna tadorna* | 1 |
| *Thalasseus sandvicensis* | 2 |
| *Trachurus trachurus* | 2 |
| *Tringa nebularia* | 1 |
| *Tringa totanus* | 1 |
| *Trisopterus esmarkii* | 2 |
| *Trisopterus luscus* | 1 |
| *Trisopterus minutus* | 1 |
| *Triturus cristatus* | 5 |
| *Turdus philomelos* | 1 |
| *Tursiops truncatus* | 4 |
| *Uria aalge* | 10 |
| *Vanellus vanellus* | 3 |
| *Xiphias gladius* | 1 |

**Table S6. Profiling method for estimating habitat specificity for 144 species with populations in the UK in the LPD.** We extracted the habitats in which each species occurs from their IUCN Red List profiles (http://www.iucnredlist.org/) and we followed this key for consistency.

| Habitat | Considered to be the same as: | Considered to be different to: |
|---|---|---|
| **Rural park** | Suburban park, urban park, rural garden, suburban garden, urban garden | |
| **Lake** | Big lake, small lake, pond, pool, dam, oxbow lake, reservoir | |
| **Bog** | Swamp, bogland | Lagoon |
| **Coastal cliff** | Island cliff | |
| **Shingle beach** | Pebble beach, rock beach | Sandy beach |
| **Stream** | River | Weir |
| **Fruit Tree Plantation** | Fruit Garden, orchard | |
| **Thicket** | Copse, grove, small stand | |
| **Forest** | | Woodland |
| **Glade** | Forest Clearing | |
| **Broadleaf** | Deciduous | |
| **Urban** | Suburban | |
| **River margin** | Various types of river margins | |
| **Tidal Creek** | | Estuary |
| **Harbour** | Dock, jetty, pier | |
| **Bush lands** | Shrublands | |
| **Irrigation channel** | Ditch | |
| **Heath** | Moorland | |
| **Sandy beach** | Spit, dune | Shingle beach, pebble beach, rock beach |
| **Crag** | Rocky outcrop, cliff, rocky slope | |
| **Marsh** | Wet meadow | |
| **Islet** | Island | |

**Table S7. References for eighty time series (or 1% of analysed time series) which had very little variance (error < 0.001).** See Figure S7e for a visualisation of the data from a subsample of those time series.

| Time series id | Data source citation |
|---|---|
| 4178 | Environment Canada (2015). North American Breeding Bird Survey - Canadian Trends Website. Data-version 2014. from http://www.ec.gc.ca/ron-bbs/P001/A001/?lang=e. |
| 13773 | Environment Canada (2015). North American Breeding Bird Survey - Canadian Trends Website. Data-version 2014. from http://www.ec.gc.ca/ron-bbs/P001/A001/?lang=e. |
| 3697 | Environment Canada (2015). North American Breeding Bird Survey - Canadian Trends Website. Data-version 2014. from http://www.ec.gc.ca/ron-bbs/P001/A001/?lang=e. |
| 18118 | Environment Canada (2015). North American Breeding Bird Survey - Canadian Trends Website. Data-version 2014. from http://www.ec.gc.ca/ron-bbs/P001/A001/?lang=e. |
| 13797 | Environment Canada (2015). North American Breeding Bird Survey - Canadian Trends Website. Data-version 2014. from http://www.ec.gc.ca/ron-bbs/P001/A001/?lang=e. |
| 4218 | Environment Canada (2015). North American Breeding Bird Survey - Canadian Trends Website. Data-version 2014. from http://www.ec.gc.ca/ron-bbs/P001/A001/?lang=e. |
| 4220 | Environment Canada (2015). North American Breeding Bird Survey - Canadian Trends Website. Data-version 2014. from http://www.ec.gc.ca/ron-bbs/P001/A001/?lang=e. |
| 3725 | Environment Canada (2015). North American Breeding Bird Survey - Canadian Trends Website. Data-version 2014. from http://www.ec.gc.ca/ron-bbs/P001/A001/?lang=e. |
| 2916 | Sauer, J. R., J. E. Hines, et al. (2012). The North American Breeding Bird Survey, Resuts and Analysis 1966 - 2011, USGS Patuxent Wildlife Research Center, Laurel, MD. |
| 13846 | Environment Canada (2015). North American Breeding Bird Survey - Canadian Trends Website. Data-version 2014. from http://www.ec.gc.ca/ron-bbs/P001/A001/?lang=e. |
| 3771 | Environment Canada (2015). North American Breeding Bird Survey - Canadian Trends Website. Data-version 2014. from http://www.ec.gc.ca/ron-bbs/P001/A001/?lang=e. |
| 2987 | Sauer, J. R., J. E. Hines, et al. (2012). The North American Breeding Bird Survey, Resuts and Analysis 1966 - 2011, USGS Patuxent Wildlife Research Center, Laurel, MD. |
| 4236 | Environment Canada (2015). North American Breeding Bird Survey - Canadian Trends Website. Data-version 2014. from http://www.ec.gc.ca/ron-bbs/P001/A001/?lang=e. |
| 4237 | Environment Canada (2015). North American Breeding Bird Survey - Canadian Trends Website. Data-version 2014. from http://www.ec.gc.ca/ron-bbs/P001/A001/?lang=e. |
| 3803 | Environment Canada (2015). North American Breeding Bird Survey - Canadian Trends Website. Data-version 2014. from http://www.ec.gc.ca/ron-bbs/P001/A001/?lang=e. |
| 3808 | Environment Canada (2015). North American Breeding Bird Survey - Canadian Trends Website. Data-version 2014. from http://www.ec.gc.ca/ron-bbs/P001/A001/?lang=e. |
| 3816 | Environment Canada (2015). North American Breeding Bird Survey - Canadian Trends Website. Data-version 2014. from http://www.ec.gc.ca/ron-bbs/P001/A001/?lang=e. |
| 3813 | Environment Canada (2015). North American Breeding Bird Survey - Canadian Trends Website. Data-version 2014. from http://www.ec.gc.ca/ron-bbs/P001/A001/?lang=e. |

| | |
|---|---|
| 3821 | Environment Canada (2015). North American Breeding Bird Survey - Canadian Trends Website. Data-version 2014. from http://www.ec.gc.ca/ron-bbs/P001/A001/?lang=e. |
| 13944 | Environment Canada (2015). North American Breeding Bird Survey - Canadian Trends Website. Data-version 2014. from http://www.ec.gc.ca/ron-bbs/P001/A001/?lang=e. |
| 11350 | KeiÅÁs, O. (2005). Impact of changes in agricultural land use on the Corncrake Crex crex population in Latvia. Acta Universitatis Latviensis 691: 93-109. |
| 13976 | Environment Canada (2015). North American Breeding Bird Survey - Canadian Trends Website. Data-version 2014. from http://www.ec.gc.ca/ron-bbs/P001/A001/?lang=e. |
| 3997 | Environment Canada (2015). North American Breeding Bird Survey - Canadian Trends Website. Data-version 2014. from http://www.ec.gc.ca/ron-bbs/P001/A001/?lang=e. |
| 3995 | Environment Canada (2015). North American Breeding Bird Survey - Canadian Trends Website. Data-version 2014. from http://www.ec.gc.ca/ron-bbs/P001/A001/?lang=e. |
| 3998 | Environment Canada (2014). North American Breeding Bird Survey - Canadian Trends Website. Data-version 2012. from http://www.ec.gc.ca/ron-bbs/P001/A001/?lang=e. |
| 13694 | Environment Canada (2015). North American Breeding Bird Survey - Canadian Trends Website. Data-version 2014. from http://www.ec.gc.ca/ron-bbs/P001/A001/?lang=e. |
| 13520 | Environment Canada (2015). North American Breeding Bird Survey - Canadian Trends Website. Data-version 2014. from http://www.ec.gc.ca/ron-bbs/P001/A001/?lang=e. |
| 3018 | Sauer, J. R., J. E. Hines, et al. (2012). The North American Breeding Bird Survey, Results and Analysis 1966 - 2011, USGS Patuxent Wildlife Research Center, Laurel, MD. |
| 13687 | Environment Canada (2015). North American Breeding Bird Survey - Canadian Trends Website. Data-version 2014. from http://www.ec.gc.ca/ron-bbs/P001/A001/?lang=e. |
| 14076 | Environment Canada (2015). North American Breeding Bird Survey - Canadian Trends Website. Data-version 2014. from http://www.ec.gc.ca/ron-bbs/P001/A001/?lang=e. |
| 14074 | Environment Canada (2015). North American Breeding Bird Survey - Canadian Trends Website. Data-version 2014. from http://www.ec.gc.ca/ron-bbs/P001/A001/?lang=e. |
| 3326 | Environment Canada (2015). North American Breeding Bird Survey - Canadian Trends Website. Data-version 2014. from http://www.ec.gc.ca/ron-bbs/P001/A001/?lang=e. |
| 14095 | Environment Canada (2015). North American Breeding Bird Survey - Canadian Trends Website. Data-version 2014. from http://www.ec.gc.ca/ron-bbs/P001/A001/?lang=e. |
| 14093 | Environment Canada (2015). North American Breeding Bird Survey - Canadian Trends Website. Data-version 2014. from http://www.ec.gc.ca/ron-bbs/P001/A001/?lang=e. |
| 13584 | Environment Canada (2015). North American Breeding Bird Survey - Canadian Trends Website. Data-version 2014. from http://www.ec.gc.ca/ron-bbs/P001/A001/?lang=e. |
| 3883 | Environment Canada (2015). North American Breeding Bird Survey - Canadian Trends Website. Data-version 2014. from http://www.ec.gc.ca/ron-bbs/P001/A001/?lang=e. |
| 10449 | Tofft, J. (2007). Tranens Grus grus bestandsudvikling i Danmark 1990-2006. Dansk Ornitologisk Forenings Tidsskrift 101(4): 67-72. |
| 2579 | Fylkesmannen i Vestfold (2004). Hekketakseringer, sjÌüfugl i Vestfold, MiljÌüvernavdelingen. |
| 2632 | Fylkesmannen i Vestfold (2004). Hekketakseringer, sjÌüfugl i Vestfold, MiljÌüvernavdelingen. |
| 3360 | Environment Canada (2015). North American Breeding Bird Survey - Canadian Trends Website. Data-version 2014. from http://www.ec.gc.ca/ron-bbs/P001/A001/?lang=e. |

| | |
|---|---|
| 14286 | Environment Canada (2015). North American Breeding Bird Survey - Canadian Trends Website. Data-version 2014. from http://www.ec.gc.ca/ron-bbs/P001/A001/?lang=e. |
| 4245 | Environment Canada (2014). North American Breeding Bird Survey - Canadian Trends Website. Data-version 2012. from http://www.ec.gc.ca/ron-bbs/P001/A001/?lang=e. |
| 4247 | Environment Canada (2015). North American Breeding Bird Survey - Canadian Trends Website. Data-version 2014. from http://www.ec.gc.ca/ron-bbs/P001/A001/?lang=e. |
| 18111 | Environment Canada (2015). North American Breeding Bird Survey - Canadian Trends Website. Data-version 2014. from http://www.ec.gc.ca/ron-bbs/P001/A001/?lang=e. |
| 4229 | Environment Canada (2015). North American Breeding Bird Survey - Canadian Trends Website. Data-version 2014. from http://www.ec.gc.ca/ron-bbs/P001/A001/?lang=e. |
| 1037 | Environment Canada (2015). North American Breeding Bird Survey - Canadian Trends Website. Data-version 2014. from http://www.ec.gc.ca/ron-bbs/P001/A001/?lang=e. |
| 14728 | Environment Canada (2015). North American Breeding Bird Survey - Canadian Trends Website. Data-version 2014. from http://www.ec.gc.ca/ron-bbs/P001/A001/?lang=e. |
| 14208 | Environment Canada (2015). North American Breeding Bird Survey - Canadian Trends Website. Data-version 2014. from http://www.ec.gc.ca/ron-bbs/P001/A001/?lang=e. |
| 3947 | Environment Canada (2015). North American Breeding Bird Survey - Canadian Trends Website. Data-version 2014. from http://www.ec.gc.ca/ron-bbs/P001/A001/?lang=e. |
| 4254 | Environment Canada (2015). North American Breeding Bird Survey - Canadian Trends Website. Data-version 2014. from http://www.ec.gc.ca/ron-bbs/P001/A001/?lang=e. |
| 4253 | Environment Canada (2015). North American Breeding Bird Survey - Canadian Trends Website. Data-version 2014. from http://www.ec.gc.ca/ron-bbs/P001/A001/?lang=e. |
| 14217 | Environment Canada (2014). North American Breeding Bird Survey - Canadian Trends Website. Data-version 2012. from http://www.ec.gc.ca/ron-bbs/P001/A001/?lang=e. |
| 3958 | Environment Canada (2015). North American Breeding Bird Survey - Canadian Trends Website. Data-version 2014. from http://www.ec.gc.ca/ron-bbs/P001/A001/?lang=e. |
| 3245 | Sauer, J. R., J. E. Hines, et al. (2012). The North American Breeding Bird Survey, Results and Analysis 1966 - 2011, USGS Patuxent Wildlife Research Center, Laurel, MD. |
| 14249 | Environment Canada (2015). North American Breeding Bird Survey - Canadian Trends Website. Data-version 2014. from http://www.ec.gc.ca/ron-bbs/P001/A001/?lang=e. |
| 6515 | Herrero, M. A. N. (2006). Results of a 10-years ( 1994-2003) monitoring Programme of Shore Birds Populations in the Protected landscape of Rambla Salada and Ajauque ( Inner Saltworks) in Murcia, Spain. A Contribution for 2010 Biodiversity Index. |
| 4261 | Environment Canada (2014). North American Breeding Bird Survey - Canadian Trends Website. Data-version 2012. from http://www.ec.gc.ca/ron-bbs/P001/A001/?lang=e. |
| 14327 | Environment Canada (2015). North American Breeding Bird Survey - Canadian Trends Website. Data-version 2014. from http://www.ec.gc.ca/ron-bbs/P001/A001/?lang=e. |
| 3674 | Environment Canada (2015). North American Breeding Bird Survey - Canadian Trends Website. Data-version 2014. from http://www.ec.gc.ca/ron-bbs/P001/A001/?lang=e. |
| 13636 | Environment Canada (2015). North American Breeding Bird Survey - Canadian Trends Website. Data-version 2014. from http://www.ec.gc.ca/ron-bbs/P001/A001/?lang=e. |

| | |
|---|---|
| 4051 | Environment Canada (2015). North American Breeding Bird Survey - Canadian Trends Website. Data-version 2014. from http://www.ec.gc.ca/ron-bbs/P001/A001/?lang=e. |
| 4069 | Environment Canada (2015). North American Breeding Bird Survey - Canadian Trends Website. Data-version 2014. from http://www.ec.gc.ca/ron-bbs/P001/A001/?lang=e. |
| 4086 | Environment Canada (2015). North American Breeding Bird Survey - Canadian Trends Website. Data-version 2014. from http://www.ec.gc.ca/ron-bbs/P001/A001/?lang=e. |
| 4088 | Environment Canada (2015). North American Breeding Bird Survey - Canadian Trends Website. Data-version 2014. from http://www.ec.gc.ca/ron-bbs/P001/A001/?lang=e. |
| 4083 | Environment Canada (2015). North American Breeding Bird Survey - Canadian Trends Website. Data-version 2014. from http://www.ec.gc.ca/ron-bbs/P001/A001/?lang=e. |
| 3119 | Sauer, J. R., J. E. Hines, et al. (2012). The North American Breeding Bird Survey, Results and Analysis 1966 - 2011, USGS Patuxent Wildlife Research Center, Laurel, MD. |
| 4085 | Environment Canada (2015). North American Breeding Bird Survey - Canadian Trends Website. Data-version 2014. from http://www.ec.gc.ca/ron-bbs/P001/A001/?lang=e. |
| 4087 | Environment Canada (2015). North American Breeding Bird Survey - Canadian Trends Website. Data-version 2014. from http://www.ec.gc.ca/ron-bbs/P001/A001/?lang=e. |
| 4093 | Environment Canada (2015). North American Breeding Bird Survey - Canadian Trends Website. Data-version 2014. from http://www.ec.gc.ca/ron-bbs/P001/A001/?lang=e. |
| 5700 | Bailey, K. M. and S. A. Macklin (1994). Analysis of patterns in larval walleye pollpck Theragra chalcogramma survival and wind mixing events in Shelikof Strait Gulf of Alaska. Marine Ecology Progress Series 113: 1-12. |
| 4099 | Environment Canada (2015). North American Breeding Bird Survey - Canadian Trends Website. Data-version 2014. from http://www.ec.gc.ca/ron-bbs/P001/A001/?lang=e. |
| 8476 | Lee, P.-F., I. C. Chen, et al. (2005). Spatial and temporal distribution patterns of bigeye tuna (Thunnus obesus) in the Indian Ocean. Zoological Studies 44(2): 260-270. |
| 14807 | Environment Canada (2015). North American Breeding Bird Survey - Canadian Trends Website. Data-version 2014. from http://www.ec.gc.ca/ron-bbs/P001/A001/?lang=e. |
| 14801 | Environment Canada (2015). North American Breeding Bird Survey - Canadian Trends Website. Data-version 2014. from http://www.ec.gc.ca/ron-bbs/P001/A001/?lang=e. |
| 14804 | Environment Canada (2015). North American Breeding Bird Survey - Canadian Trends Website. Data-version 2014. from http://www.ec.gc.ca/ron-bbs/P001/A001/?lang=e. |
| 4132 | Environment Canada (2015). North American Breeding Bird Survey - Canadian Trends Website. Data-version 2014. from http://www.ec.gc.ca/ron-bbs/P001/A001/?lang=e. |
| 4143 | Environment Canada (2015). North American Breeding Bird Survey - Canadian Trends Website. Data-version 2014. from http://www.ec.gc.ca/ron-bbs/P001/A001/?lang=e. |
| 4139 | Environment Canada (2015). North American Breeding Bird Survey - Canadian Trends Website. Data-version 2014. from http://www.ec.gc.ca/ron-bbs/P001/A001/?lang=e. |
| 4151 | Environment Canada (2015). North American Breeding Bird Survey - Canadian Trends Website. Data-version 2014. from http://www.ec.gc.ca/ron-bbs/P001/A001/?lang=e. |
| 18190 | Environment Canada (2015). North American Breeding Bird Survey - Canadian Trends Website. Data-version 2014. from http://www.ec.gc.ca/ron-bbs/P001/A001/?lang=e. |

241

**Table S8. References for time series which appear to show logistic growth with little variance (see Figure S7e for visualisation of data).**

| Time series id | Data source citation |
|---|---|
| 468 | NERC Centre for Population Biology (1999). The Global Populations Dynamics Database. http://cpbnts1.bio.ic.ac.uk/gpdd/, Imperial College,.l Batten, L. A. and J. H. Marchant (1977). Bird Population Changes for Years 1974-75. Bird Study 24(1): 55-61. |
| 17803 | Government of Antigua and Barbuda (2014). Antigua and Barbuda Fifth National Report to the Convention on Biodiversity, Environment Division: 1-66. |
| 10193 | Giling, D., R. D. Reina, et al. (2008). Anthropogenic influence on an urban colony of the little penguin Eudyptula minor. Marine and Freshwater Research 59(7): 647-651. |
